# Structural basis for a phosphoinositide-driven mTORC2-AKT positive feedback loop

**DOI:** 10.64898/2026.01.08.698367

**Authors:** Iain M. Hay, Maxime Bourguet, Bilal Ahsan, Olga Perisic, Madhangopal Anandapadamanaban, Roger L. Williams

## Abstract

The mammalian target of rapamycin complex 2 (mTORC2) regulates metabolism, growth, survival and cytoskeletal organization, yet its activation mechanism is poorly understood. We show that mTORC2 is directly activated by membranes and our cryo-electron tomography structure of membrane-bound mTORC2 reveals the N-HEAT region of mTOR is at the major membrane interface. mTORC2 is further potently activated by a positive feedback loop involving reciprocal phosphorylation of mTORC2 and its substrate kinase AKT. Cryo-EM structures of dephosphorylated, autophosphorylated and AKT-phosphorylated mTORC2 reveal structural changes in the SIN1 subunit, regulating an autoinhibitory anchor. Reconstitution of the PDK1–AKT–mTORC2 hub on PIP3-containing membranes shows that PDK1/PIP3-dependent AKT activation drives SIN1-T86 phosphorylation, enabling mTORC2 to phosphorylate S473 of AKT’s hydrophobic motif, establishing a PI3K-dependent, phosphorylation-driven positive feedback loop at the membrane.

## Main text

The serine/threonine protein kinase mammalian target of rapamycin (mTOR) is a key regulator of homeostasis and development, integrating signaling from growth factors and nutrient availability to coordinate cell metabolism, growth, survival and proliferation. The mTOR catalytic subunit is the central component of two distinct multi-subunit complexes, mTOR complex 1 (mTORC1) and mTORC2. In contrast to mTORC1, which has extensively studied regulation by nutrients and growth factors, mTORC2 regulation is less well understood. mTORC2 participates in both anabolic and stress-adaptive processes, phosphorylating and activating AGC kinases, notably AKT, SGK, PKC and PKN, to promote survival, proliferation, and cytoskeletal organization (*1–4*). mTORC2 activation is closely associated with

PI3K signaling and is regulated by growth factors and nutrients, but sensitive to contexts such as sub-cellular localization, which makes its regulation more complicated than mTORC1 (*3, 5*).

mTORC2 has a critical role in the function of the master kinase AKT. AKT is partially activated by PDK1 phosphorylating the AKT activation loop at T308, but fully active AKT also requires phosphorylation in its hydrophobic motif (HM) at S473 (*6, 7*). The HM of AKT is directly phosphorylated by mTORC2 (*8, 9*). However, a different model was recently proposed, in which mTORC2 phosphorylates AKT on T443 in a TOR interaction motif (TIM), which, in turn, facilitates AKT autophosphorylation of S473 (*10*). Activated pT308/pS473-AKT can stimulate both anabolic mTORC1-dependent pathways as well as inhibit FOXO3/1 transcription factors to promote cell survival (*11*).

mTORC2 is localized to several membrane compartments, including plasma membranes, endosomal vesicles, and mitochondria (comprehensively reviewed in (*5*)), but what brings it there and the impact of localization on its activity and regulation remains highly enigmatic (*1, 5*). At the plasma membrane, mTORC2 activity is PI3K independent, whereas on endosomal membranes the activity is PI3K dependent (*12*). mTORC2 physically associates with assembled and actively translating ribosomes (*1, 13, 14*). The interaction with ribosomes is linked to the abundant presence of mTORC2 on ER and mitochondria-associated ER membranes (MAMs) (*15, 16*). This ribosomal association activates mTORC2 and enables it to co-translationally phosphorylate the AKT C-terminal tail at T450 in a region known as the turn motif (TM), which serves as a quality-control mechanism for Akt translation. When T450 phosphorylation is ablated, Akt becomes ubiquitylated and is degraded (*13, 17*).

Structurally, mTORC2 is a dimer of protomers related by two-fold rotational symmetry. Each mTORC2 protomer is a heterotetramer consisting of mTOR, mLST8, rapamycin-insensitive companion of mammalian TOR (RICTOR) and stress-activated map kinase-interacting protein 1 (SIN1). SIN1, RICTOR and mLST8 subunits are essential for mTORC2 function, and mice with RICTOR, mLST8 or SIN1 ablated die at E10.5-E11.5 (*18–20*). Within the N-lobe of the mTOR kinase domain, there is an insertion of a helical bundle known as the FRB domain (fragment for rapamycin binding) that binds the rapamycin/FKBP12 complex. The insensitivity of mTORC2 to rapamycin stems from C-terminal domain of RICTOR/Avo3 subunit occluding its FRB-binding site (*21, 22*).

The SIN1 subunit of mTORC2 has several different isoforms, generated by alternative splicing, with the longest isoform containing four regions: the N-terminal region (NTD), the conserved region in the middle (CRIM), the Ras-binding domain (RBD) and the pleckstrin homology (PH) domain (*23*). Only splice variants that contain the N-terminal region assemble into mTORC2 (*24*). The high-resolution cryo-EM structure of mTORC2 showed that the NTD of SIN1 interacts with RICTOR and mLST8 subunits (*21*). The CRIM domain of SIN1 is responsible for recruiting mTORC2 substrate kinases, with the acidic loop protruding from the CRIM domain playing a critical role in the binding to AGC substrate kinases (*25–27*). Very recently, a tour-de force cryo-EM reconstruction of mTORC2 with a bisubstrate AKT1-Torin1 analog revealed bipartite engagement of the AKT1 kinase domain with mTORC2, with the N-lobe of AKT1 interacting with the CRIM domain of SIN1, and the C-lobe of AKT1 interacting with the N-terminal domain of SIN1 (*28*). In contrast to the clear role of the SIN1 CRIM domain, the role of the SIN1 PH domain remains enigmatic. The PH domain of SIN1 is not visible in the cryo-EM reconstructions of mTORC2, suggesting flexible attachment, although pull-down studies of truncation variants suggested that the SIN1 PH domain makes an inhibitory interaction with the mTORC2 kinase domain (*29*). It was proposed that the interaction of SIN1 PH domain with PIP3 breaks this autoinhibitory SIN1/mTOR interaction, leading to PIP3-dependent activation of mTORC2 (*29*). However, another study showed that SIN1 and mTORC2 localization to the plasma membrane is independent of PI3K (*12*), consistent with observations that the isolated SIN1/Avo1 PH domain binds phosphatidylinositol and phosphorylated phosphoinositides but is not specific to PI3K products (*23, 30, 31*). A recent structural study of yeast TOR2 in the TORC2 complex lends support for an autoinhibitory role of the PH domain of the yeast SIN1 orthologue, Avo1, with the PH domain contacting the TOR kinase active site. This role is not simple, since deletion of the SIN1 PH domain resulted in less TORC2 activity (*32*). In mammalian mTORC2, deletion of the mSIN1 PH domain decreased mTORC2 plasma membrane localization and activity in cells (*12, 24, 28*).

Although SIN1 has an essential role in mTORC2 function, its role in fine-tuning mTORC2 activity has been controversial. Phosphorylation of SIN1 at T86 and T398 by S6K was reported to prevent SIN1 from incorporating into mTORC2, thereby inhibiting mTORC2 (*33*). However, other studies showed that AKT phosphorylates SIN1-T86 to increase mTORC2 activity and that mTORC2 and AKT are involved in a positive feedback loop (*34, 35*). Cryo-EM analysis showed that Sin1-T86 is part of an interaction with the RICTOR subunit, with T86 binding in a negatively charged pocket on the RICTOR surface (*21*). This suggested that phosphorylation of T86 might diminish the interaction with RICTOR, leading to destabilization of the mTORC2 complex, which would be consistent with the previous report of mTORC2 inhibition by pT86.

The molecular sequence of events that recruit and activate mTORC2 remain important areas of investigation. mTORC2 localizes to cellular membranes, but the structural determinants of membrane binding are unknown. While the role of mTORC2 in activating AKT is widely accepted, the consequence of SIN1 phosphorylation by AKT (whether mTORC2 is activated or inhibited by AKT phosphorylation) remains contentious, and the corresponding structural mechanism for this regulation is unknown. Here, we combine structural and biochemical approaches to define the molecular determinants of mTORC2 activation, providing mechanistic insight into this underexplored complex. Our electron cryotomography (cryo-ET) showed that mTORC2 associates with membranes, primarily through the N-terminal mTOR N-HEAT, and functional analysis of mTORC2 on membranes showed that membranes activate mTORC2 in a PIP2-dependent manner, even in the absence of the SIN1 PH domain. Our reconstituted system directly established that there is a feedforward loop consisting of pT308-AKT phosphorylating mTORC2 SIN1-T86 to activate mTORC2, which then phosphorylates AKT-S473. We have determined the structural basis for mTORC2 activation by AKT-dependent phosphorylation from high resolution cryo-EM structures of dephosphorylated-mTORC2, autophosphorylated mTORC2, and AKT-phosphorylated mTORC2. Our data provide a mechanistic view of PDK1/AKT driven mTORC2 activation on phosphoinositide membranes.

## Results

### Activation of mTORC2 by PIP containing membranes

We reconstituted mTORC2 activity *in vitro*, using recombinant mTORC2 purified from mammalian cells. Full-length recombinant, dephosphorylated, kinase-dead (D274A) AKT1 (AKT^D274A^) was used as a model mTORC2 substrate. To assay mTORC2 activity, substrate phosphorylation was visualized by band shift on Coomassie-stained Phos-Tag acrylamide gels (**fig. S1**).

To investigate whether membrane localization may modulate mTORC2 activity, we reconstituted mTORC2 membrane binding *in vitro* using 100 nm large unilamellar vesicles (LUVs). LUVs consisted of 35% DOPE, 30% DOPC, 15% DOPS, 20% cholesterol +/- 5% phosphatidylinositol 4,5-bisphosphate (PIP2) or phosphatidylinositol 3,4,5-trisphoaphate (PIP3) (**Table S1**). We first determined the binding specificity of mTORC2 for PIP containing membranes. Liposome flotation assays on sucrose gradients showed that while mTORC2 has residual binding to membranes in the absence of PIP species, membrane recruitment was significantly enriched in both 5% PIP2 and PIP3 containing liposomes (**Fig. 1A**). Surprisingly, we found the mTORC2 SIN1-PH domain to be largely dispensable for membrane interaction, as deletion of the SIN1-PH domain within mTORC2 (mTORC2ΔPH) did not significantly alter membrane binding to either PIP2 or PIP3 vesicles (**fig. S2A and S2B**). However, flotation assays using the isolated SIN1-PH domain showed that this domain was sufficient for binding to both PIP2 and PIP3 membranes (**fig. S2C**), suggesting that while the SIN1-PH is capable of PIP binding it is not a strict requirement for membrane recruitment in the context of the intact mTORC2 complex. Strikingly, PIP2-containing LUVs were able to activate mTORC2 *in vitro*, resulting in ∼2-fold increase in enzymatic turnover compared to mTORC2 in the absence of liposomes (**Fig. 1B**). These results show that mTORC2 can bind to both PIP2 and PIP3 containing membranes, leading to mTORC2 activation.

**Fig. 1.**
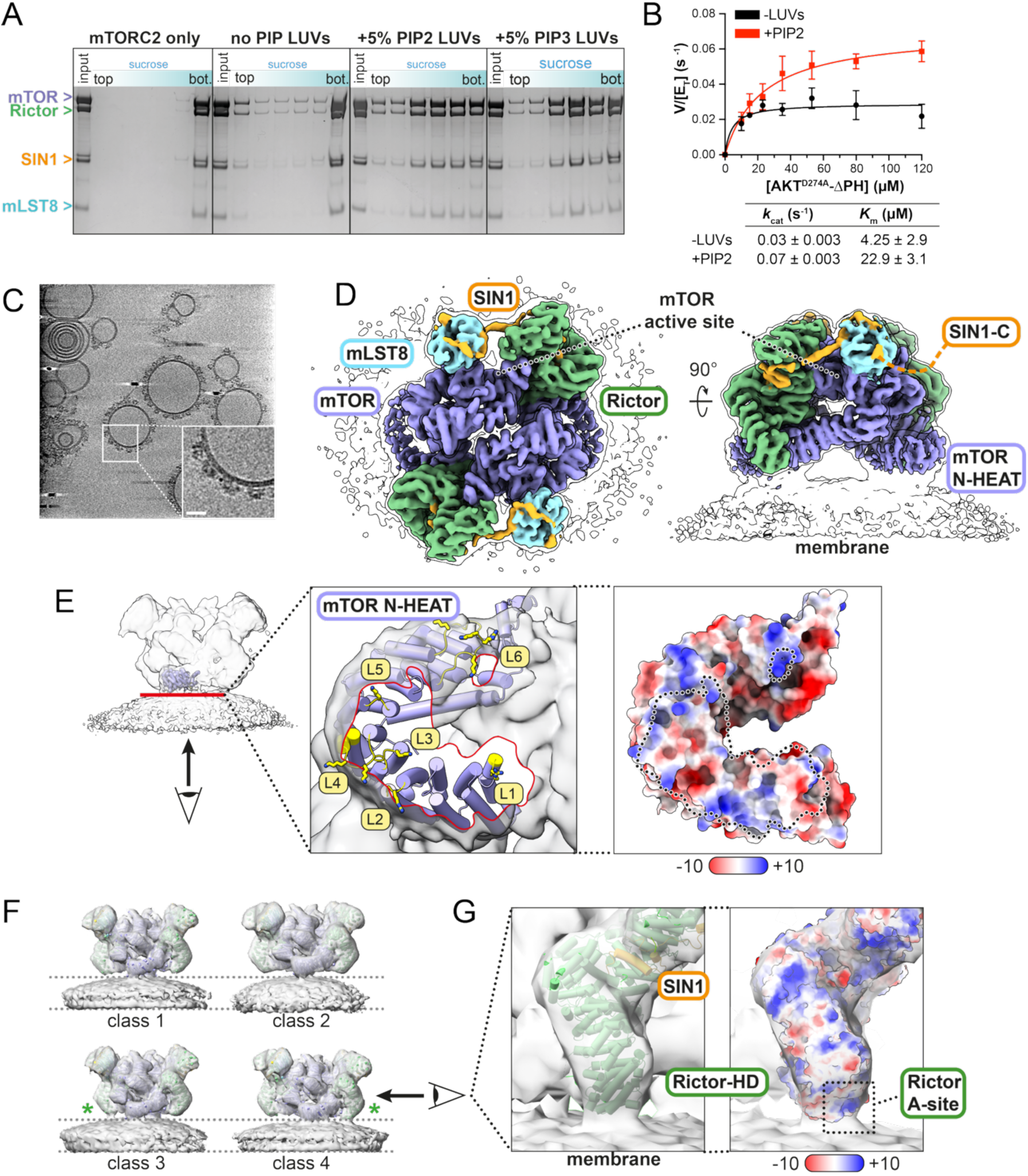
Structure and activation of membrane-bound mTORC2. (A) Liposome flotation assays of mTORC2 alone or in the presence of no PIP (base lipids only, see Table S1), 5% PIP2 or 5% PIP3 containing 100 nm LUVs in a 35% (bottom) to 25% (top) sucrose gradient. The mTORC2/lipid mixture was placed at the bottom of the gradient, with LUVs and LUV-bound proteins floating into top fractions during centrifugation. (B) Kinetic analysis of AKT^D274A^ΔPH substrate phosphorylation by mTORC2 alone (0.3 µM, black) and in the presence of 5% PIP2 containing 100 nm LUVs. Kinetic parameters are outlined in the table and data represents the mean of n = 2 independent experiments ± SEM. (C) Representative Z-slice of a reconstructed tomogram showing mTORC2 bound to PIP2 containing liposomes. Inset: Zoomed view of membrane bound mTORC2 particles, scale bar = 25 nm. (D) Cryo-ET density map of membrane-bound mTORC2. The consensus mTORC2 map is colored by each subunit as indicated and is overlaid with a low-pass filtered version of the map (black outline) to aid visualization of orientation towards the membrane. (E) Cross section of the mTOR N-HEAT membrane contact site. The Z-slice is shown (red line) on the low-pass filtered mTORC2-membrane map, with the inset showing the view of the contact area up and through the membrane. Inset left: the mTORC2 N-HEAT atomic model is show in cartoon, with membrane proximal loops colored yellow. Basic residues (Lys/Arg) within these loops are shown in stick representation. The membrane contact area is highlighted (red outline). Loops are numbered from N to C, as shown in fig. S4. Inset, right: electrostatic potential of the mTOR N-HEAT at the membrane contact site (dashed outline). (F) Four mTORC2 membrane-bound classes from 3D classification in bin4. Classes with apparent Rictor-membrane contact sites are marked with a green asterisk. (G) Left: The Rictor atomic model is shown in cartoon representation and docked into the density for class 4 from (F). Right: Electrostatic potential of Rictor, highlighting the location of the positively charged A-site within the Rictor HEAT domain (HD) proximal to the membrane.

### Structural overview of membrane-bound mTORC2

As PIP2 is the most abundant PIP species within the plasma membrane (*36, 37*) and PIP2 was sufficient to both bind and activate mTORC2, we determined a cryo-electron tomography (cryo-ET) reconstruction of mTORC2 on LUVs containing 5% PIP2 (**Fig. 1C**). A total of 38,519 sub-tomogram particles picked from 180 collected tilt-series were included in the final reconstruction, resulting in a 7.7 Å consensus map including both mTORC2 and the membrane. This could be further improved to 6.4 Å by refinement focused on the mTORC2 protein only, allowing better particle alignment by masking out the membrane (**fig. S3 and Table S2**).

Membrane-bound mTORC2 is oriented with the spiral of the mTOR N-HEAT contacting the membrane and the mTOR active site, mLST8 and SIN1 subunits oriented up and away from the membrane (**Fig. 1D**). During classification of sub-tomograms we observed no other orientation of mTORC2 on the membrane and this orientation is broadly similar as for mTOR within the recently described mTORC1-Rag/Ragulator membrane-bound complex (*38*). Our cryo-ET structure shows the major site of membrane contact is within the N-HEAT domains of the two mTOR subunits (**Fig. 1D and 1E**). This agrees with our biochemical results showing the SIN1 PH domain plays a minor role in membrane recruitment in our reconstituted system (**fig. S2**). We observed no density for the SIN1 PH-domain on the membrane in our structure. While we confirmed that the SIN1 PH domain is capable of binding PIP2 membranes (**fig. S2C**), it is likely that the SIN1-PH domain does not adopt a consistent, ordered orientation in relation to the core of the complex when engaged with the membrane and therefore would likely be averaged out during reconstruction.

Due to the inherent flexibility of the N-HEAT domain within mTOR complexes, this domain of mTOR is generally less well resolved in mTORC1 and mTORC2 structures, preventing accurate modeling. While our resolution is similarly insufficient to allow high-resolution atomic modeling, we fitted an AlphaFold3 model of the mTOR N-HEAT into our membrane-bound mTORC2 map and model. In the membrane-bound orientation, we noted several loops within the N-HEAT domain that are localized to the interface with the membrane (**Fig. 1E**). Inspection of the mTOR N-HEAT sequence (**fig. S4**) showed that many of these membrane-proximal N-HEAT loops contain residues with positively charged sidechains, suggesting charge complementarity with the negatively charged PIP2-containing membrane may stabilize mTORC2 in this specific orientation.

While the overall mTORC2 orientation with mTOR N-HEAT contacting the membrane was the only one observed in our data, during 3D classification of sub-tomograms we resolved slight variations in membrane curvature and the angle of mTORC2 relative to the membrane (**Fig. 1F**). In all classes, the N-HEAT domains in both mTOR chains are in contact with the membrane, consistent with this being the major contact site. Additionally, we observed classes in which one Rictor copy is in contact with the membrane, suggesting that the complex may ‘rock’ on an axis centered through the mTOR N-HEAT contact regions (**Fig. 1F**). This membrane-proximal region of Rictor contains the positively-charged ligand-binding ‘A-site’, that was previously observed to bind ATP (*21*) (**Fig. 1G**). While our data did not provide sufficient resolution to determine molecular interactions, like the mTOR N-HEAT loops, it is possible this positively charged Rictor patch may play a role in membrane binding via interaction with negatively charged membrane lipids such as PIP2.

### Activation of mTORC2 by AKT phosphorylation

During initial exploration of mTORC2 activity, we observed that the rates of substrate phosphorylation in mTORC2 kinase assays were higher when using WT, kinase-active AKT (AKT^WT^) versus AKT^D274A^ as mTORC2 substrates (**Fig. 2A**). Previously it was reported that AKT can phosphorylate and activate mTORC2 (*34, 35*). The role of AKT phosphorylation in mTORC2 activation remains controversial, being separately reported as either activating (*34*) or inhibiting (*33*). We therefore sought to determine whether AKT phosphorylation controls activation of mTORC2 kinase activity in our *in vitro* reconstituted system.

**Fig. 2.**
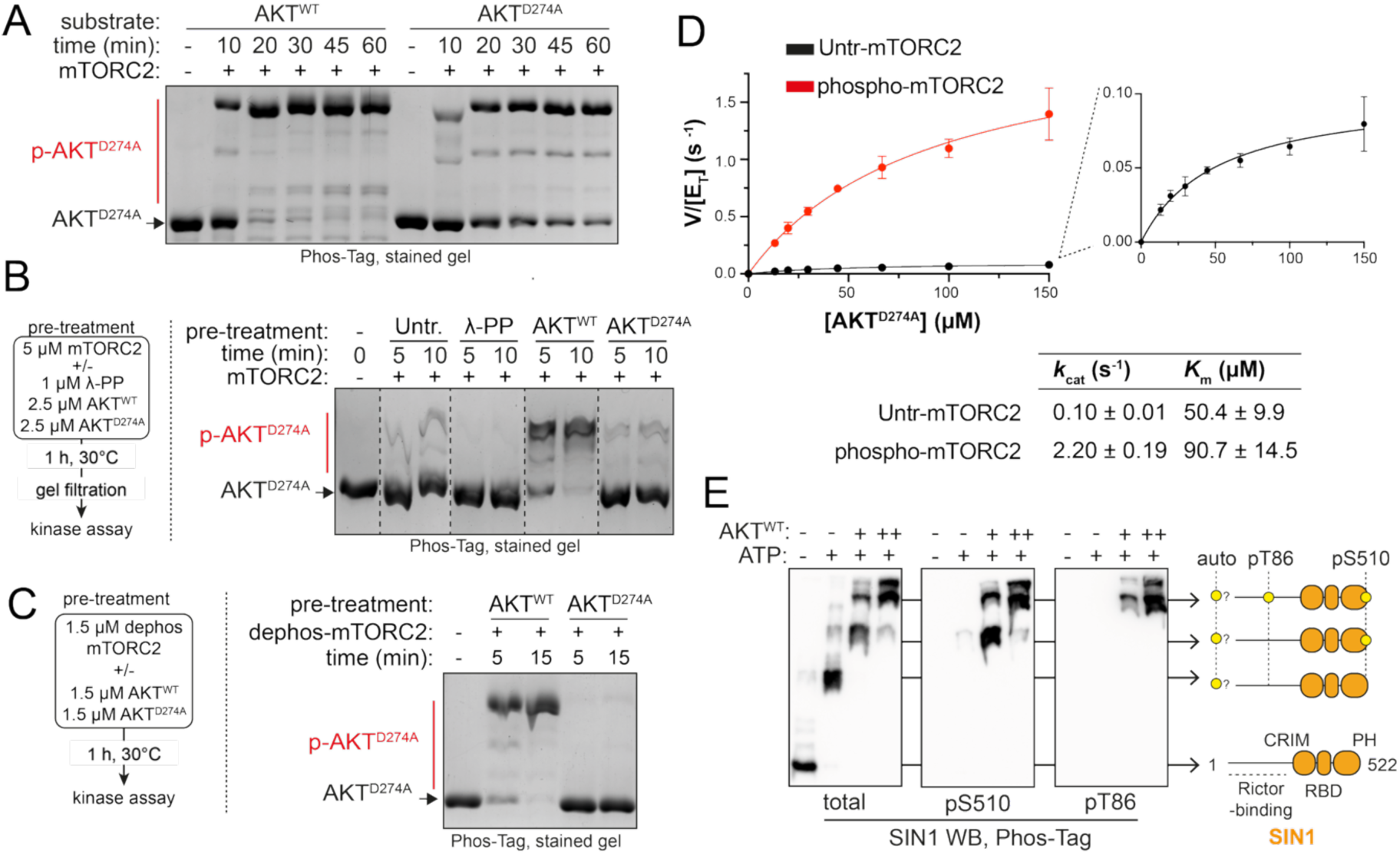
Phosphorylation by AKT activates mTORC2. (A) A time course of mTORC2 activity, using 0.3 µM mTORC2 with 25 µM AKT kinase active (WT) or kinase dead (D274A) as substrate. (B) mTORC2 activity after treatment with lambda protein phosphatase (λ-PP), AKT^WT^ or AKT^D274A^, using 0.2 µM treated mTORC2 and 25 µM AKT^D274A^ substrate. (C) mTORC2 activity assay of dephospho-mTORC2 after pre-treatment with either AKT^WT^ or AKT^D274A^, using 0.2 µM treated mTORC2 with 25 µM AKT^D274A^ substrate. (D) Kinetic analysis of AKT^D274A^ substrate phosphorylation by untreated (Untr-mTORC2, 0.3 µM, black) or AKT phosphorylated (phospho-mTORC2, 30 nM, red) mTORC2. Inset: Zoomed view of the Untr-mTORC2 data. Kinetic parameters are outlined in the table and data represents the mean of n = 3 independent experiments ± SD. (E) Analysis of SIN1 phosphorylation by Phos-Tag western blotting of 2.5 µM dephospho-mTORC2 treated with ATP alone, 0.25 µM AKT^WT^ (+) or 2.5 µM AKT^WT^ (++) for 1 h at 30°C. All phosphorylation steps/assays shown used 1 mM ATP, 10 mM MgCl2.

When mTORC2 is purified from Expi-293 mammalian cells (hereafter referred to as ‘untreated’-mTORC2) the complex retains low but detectable levels of activity *in vitro* (**Fig. 2B**). To determine if phosphorylation by AKT activates mTORC2, we incubated untreated-mTORC2 with AKT^WT^ or kinase-dead AKT^D274A^. In addition, we prepared dephosphorylated (treated) mTORC2 complex, in which we removed all basal phosphorylation of mTORC2 with lambda protein phosphatase (λ-PP). The treated mTORC2 complex was then isolated by gel filtration for use in kinase assays. AKT^WT^-treated mTORC2 showed greatly enhanced activity in comparison to both untreated and AKT^D274A^-treated mTORC2 (**Fig. 2B**). Conversely, dephosphorylation of untreated-mTORC2 with λ-PP reduced mTORC2 activity to almost undetectable levels (**Fig. 2B**). Strikingly, after inactivation by λ-PP treatment, incubation with AKT^WT^ was sufficient to reactivate the dephosphorylated mTORC2 (**Fig. 2C**). With this in mind, we generated preparative quantities of recombinant, dephosphorylated mTORC2 (dephospho-mTORC2) that was then re-phosphorylated *in vitro* with AKT^WT^ (phospho-mTORC2) (**fig. S5A and S5B**). Both dephospho- and phospho-mTORC2 complexes were stable, with similar global thermal melting profiles comparable to untreated-mTORC2 (**fig. S5C**). In agreement with previous results, untreated-mTORC2 has low baseline activity in kinetic experiments (*k*cat = 0.1 s^-1^), while phospho-mTORC2 showed an over 20-fold activation of *k*cat (2.2 s^-1^) (**Fig. 2D and fig. S5D**). Kinetic parameters for dephospho-mTORC2 could not be determined due to the extremely low activity of this enzyme, but end-point assays showed 1 μM dephospho-mTORC2 to have activity comparable to 0.1 μM untreated-mTORC2 (**fig. S5E**), suggesting the *k*cat of dephospho-mTORC2 to be in the range of ∼0.01 s^-1^. The fold-activation for dephospho- vs phospho-mTORC2 is therefore likely to be >200-fold. In summary, these data show that phosphorylation by AKT is both necessary and sufficient for potent activation of mTORC2 kinase activity. We next sought to determine which phosphorylated residues within mTORC2 drive AKT-mediated activation. We focused on the SIN1 subunit of mTORC2, due to previous reports of SIN1 phosphorylation by AKT and/or S6K (*33–35*). In dephospho-mTORC2, SIN1 migrates as a single band on PhosTag SDS-PAGE, consistent with no phosphorylation being present (**Fig. 2E and fig. S5F**). Upon incubation with ATP + AKT^WT^, we observed three distinct SIN1 phospho-species by western blotting on PhosTag acrylamide gels (**Fig. 2E**). Blotting with phospho-specific SIN1 antibodies showed that SIN1-T86 and S510 are phosphorylated in an AKT dependent manner within mTORC2. T86 is located within the SIN1 ‘bridge’ spanning the N-terminal Rictor and LST8 interacting regions (residues 85-103), while S510 is located at the C-terminus of the SIN1 PH domain (**Fig. 2E**). Interestingly, one

SIN1 phosphorylated species was generated when dephospho-mTORC2 was incubated with ATP only, showing that mTORC2 autophosphorylates its SIN1 subunit (**Fig. 2E**; whether this is cis- or trans autophosphorylation is unknown). In summary, these results show that AKT is a potent activator of mTORC2 activity and this may be driven by both AKT and mTORC2 (auto)phosphorylation within the mTORC2 subunit SIN1.

### mTORC2 auto- and AKT phosphorylation drive conformational changes in SIN1

To further investigate the mechanism of AKT phosphorylation in mTORC2 activation, we determined cryo-EM single-particle analysis (SPA) structures of dephosphorylated (dephospho-, λ-PP treated), autophosphorylated (autophospho-, incubated with ATP) and AKT phosphorylated (phospho-, incubated with ATP + AKT^WT^) mTORC2 (**fig. S6 and Table S3**). To ensure that all phosphorylation would be a result of either mTORC2 autophosphorylation or AKT-mediated phosphorylation, these samples were generated using λ-PP treated mTORC2 as the starting material, removing any heterogeneous phosphorylation carried through from purification. These complexes yielded 3.3, 3.1 and 3.0 Å overall resolution reconstructions of mTORC2 dimers of dephospho-, autophospho- and phospho-mTORC2 states, respectively (**fig. S7A**). Further refinement focused on the single hetero-tetrameric protomer with best local resolution resulted in improved resolutions of 3.0, 2.8 and 2.6 Å for dephospho-, auto- and phospho-mTORC2, respectively and these maps were used for model building (**fig. S7B**).

The phospho-mTORC2 structure was determined using a kinase reaction consisting of 1.5 µM dephospho-mTORC2, 1 µM AKT^WT^, 1 mM ATP, and 10 mM MgCl2. No density for AKT^WT^ was observed, consistent with the proposed AKT binding site for mTORC2 being in the flexibly tethered SIN1 CRIM domain (*21, 39*). We found nucleotides bound within phospho-mTORC2, with ATP bound in the mTOR active site (**fig. S8A**) and ADP bound to the previously described ‘A-site’ withing the Rictor HEAT domain (*21*) (**fig. S8B**). ADP would be generated during the kinase reaction from reciprocal mTORC2/AKT phosphorylation. While the exact ATP/ADP concentration cannot be known, as micro-molar substrate (both mTORC2 and AKT) concentrations were present with 10 mM ATP, it is likely that ATP was present in vast excess compared to ADP, suggesting the ‘A-site’ may preferentially bind ADP. Finally, we observed IP6 binding in all structures at the previously described ‘I-site’ within the mTOR FAT domain (*21*) (**fig. S8C**).

Across the three mTORC2 phosphorylation states, we observed the most significant conformational changes within the SIN1 subunit. The general topology of SIN1 is outlined in **Fig. 3A** and described as follows. The acetylated SIN1 N-terminus is buried in a groove between the Rictor ARM (26–486) and HEAT (487–1007) domains, followed by the SIN1 helical α1/α2 hairpin that is anchored onto the face of Rictor ARM domain helices α7, α9, and α12. The SIN1 hairpin is followed by the SGK-Targeting, Rictor-InteractING (string) region of SIN1 (residues 66-84), which extends across the face of Rictor helices α2, α4, and α7 in the ARM domain (*39*). Immediately downstream of the string, helix α3 in the SIN1 ‘bridge’ region (SIN1 residues 85-103, also referred to as the SIN1 ‘traverse’) spans a deep cleft above the mTOR active site, bridging the Rictor and LST8 interacting regions of SIN1 (*21*). Consistent with previous studies, the C-terminal SIN1 CRIM, RBD and PH (including the putative pS510 AKT phosphorylation site) domains are flexibly tethered and not observed in the structure (*21, 39*).

**Fig. 3.**
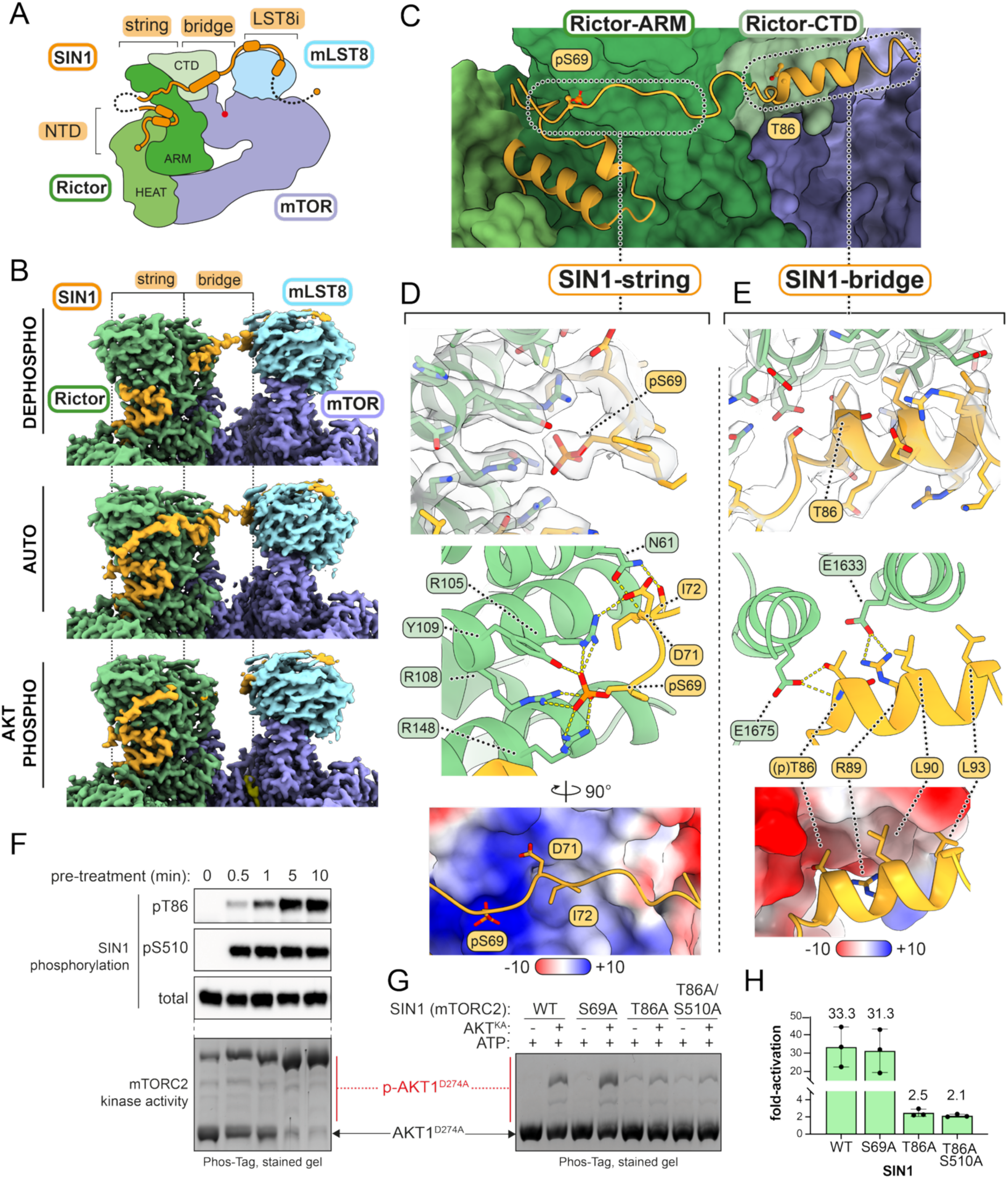
Phosphorylation of SIN1 induces structural rearrangement within mTORC2. (A) Schematic of SIN1 organization within a single heterotetramer of the mTORC2 dimer. (B) Sharpened cryo-EM density maps highlighting differential ordering of the SIN1 string and bridge regions across dephosphorylated (dephospho), autophosphorylated (auto) and AKT-phosphorylated (AKT phospho) mTORC2 datasets. The density is colored by each subunit as indicated. (C-E) Focused views of the SIN1 string-bridge (orange) interactions with Rictor (green). Key residues are shown as sticks throughout and labelled. These panels show the auto-phosphorylated mTORC2 model, as this dataset contained both ordered string and bridge SIN1 regions simultaneously. Corresponding model views for all mTORC2 phosphorylation states are shown in fig. S10. (C) Overview of SIN1 string and bridge interactions with the Rictor ARM (dark green surface) and C-terminal domains (light green surface). (D) Molecular details of SIN1 string to Rictor-ARM interaction. Top: Sharpened cryo-EM density map (transparent surface) highlighting phosphorylation of SIN1-S69. Middle: Key interactions between the Rictor ARM domain and the phosphorylated SIN1 string. Putative salt bridges (Rictor-R105, R108 and R148) and H-bonding (Rictor-Y109) to SIN1-pS69 are shown as yellow dashes. Bottom: Electrostatic potential of the Rictor-ARM surface highlighting SIN1-pS69 position within a positively charged Rictor pocket. (E) Molecular details of SIN1 bridge to Rictor-CTD interaction. Top: Sharpened cryo-EM density map (transparent surface) highlighting the position of SIN1-T86. Middle: Key interactions between the Rictor-CTD and SIN1 bridge. Putative H-bonds are shown as yellow dashes. Bottom: Electrostatic potential of the Rictor-CTD surface highlighting non-phosphorylated SIN1-T86 within a negatively charged Rictor pocket. (F) Correlation of SIN1 phosphorylation with mTORC2 activity. Equimolar mTORC2 and AKT (1 µM) were pre-phosphorylated for the indicated time and either analyzed by western blot or seeded into mTORC2 activity assays at 0.1 µM with 25 µM of AKT^D274A^ substrate for 10 min at 30°C. (G) Activation of SIN1 mutant mTORC2 by AKT^WT^. The indicated mutants (1 µM) were incubated with ATP alone (-) or with ATP + 1 µM AKT^WT^ (+) for 30 min at 30°C and seeded into mTORC2 activity assays at 0.1 µM with 25 µM AKT^D274A^ substrate for 5 min at 30°C. (H) Quantification of fold-change upon stimulation of mTORC2 mutants by AKT^WT^ versus incubation with ATP alone from (G). Data represents the mean of n = 3 independent experiments ± SD. Prephosphorylation steps and kinase assays shown utilized 1 mM ATP, 10 mM MgCl2.

We observed several key order-disorder transitions in the SIN1 string and bridge regions across the phosphorylated states of mTORC2. The SIN1-string was visible in both autophospho- and phospho-mTORC2, but not dephospho-mTORC2; whilst the SIN1-bridge was observed in dephospho- and auto-mTORC2 but not in phospho-mTORC2 (**Fig. 3B**). Ordering of the SIN1-string region in auto- (residues 66-84) and phospho-mTORC2 (residues 67-78) coincided with clear density for phosphorylation of SIN1-S69 in both structures (**Fig. 3D**). The negatively charged phosphate group of SIN1-pS69 occupies a complementary basic pocket on the face of the Rictor ARM domain, consisting of Rictor residues R105, R108, R148 and Y109 in helices α4 and α7, forming salt-bridges (Arg) and H-bonds (Tyr), with SIN1-pS69 (**Fig. 3D**). Generation of a SIN1-S69A mutant mTORC2 prevented SIN1 autophosphorylation of dephospho-mTORC2 when incubated with ATP, confirming pS69 to be the observed SIN1 autophosphorylation site (**fig. S9A**). Consistent with the structural importance of the interaction between SIN1 pS69 and the Rictor basic pocket, the entire string region is disordered in dephospho-mTORC2 (**Fig. 3B**), suggesting that autophosphorylation at SIN1-S69 anchors the string to the α2/α4/α7 face of the Rictor ARM domain.

The SIN1 bridge is observed in both dephospho- and autophospho-mTORC2 structures (**Fig. 3B and fig. S10**). In autophospho-mTORC2, residues 66-103 are ordered, with the bridge running continuously from the string region to the beginning of the LST8-interacting region. Due to string disorder in dephospho-mTORC2, only SIN1 residues 82-103 are ordered, which include the very C-terminal string residues 82-84 and the bridge (residues 85-103). In both cases, the SIN1-T86 hydroxyl and backbone amide form H-bonds with Rictor-E1675 in helix CTDα5, with further stabilizing interactions between SIN1-R89 and Rictor-E1633 in helix CTDα3 (**Fig. 3E**). SIN1-T86 within the bridge helix occupies an acidic patch on the Rictor-CTD comprising CTD α3 and α5 (**Fig. 3E**). Upon AKT phosphorylation, we observed complete disordering of the SIN1 bridge, encompassing the very C-terminal portion of the string through to the first residue of the LST8i region (residues 79-103 disordered) (**Fig. 3B and fig. S10**). Our data suggest that phosphorylation of T86 by AKT would break the SIN1-T86/Rictor-E1675 interaction due to electrostatic repulsion of pT86 from the negatively charged face of Rictor, resulting in disordering of the SIN1 bridge.

### SIN1-T86 phosphorylation is critical for mTORC2 activation

Based on these structural observations, we next sought to determine whether disordering of the SIN1 bridge via SIN1-T86 phosphorylation is a driver of mTORC2 activation by AKT. Analysis of an AKT SIN1 phosphorylation timecourse in tandem with mTORC2 activity assays showed that while both SIN1 T86 and S510 are phosphorylated by AKT during mTORC2 activation, only SIN1-pT86 correlates with activated mTORC2 (**Fig. 3F**). We generated mTORC2 with mutations in key phosphorylated residues within SIN1: S69A (autophosphorylation site), T86A and a T86A/S510A double mutant (AKT phosphorylation sites). To test which SIN1 phosphorylation sites contribute to mTORC2 activation, we pre-treated these mutant complexes with either ATP alone or ATP + AKT^WT^ prior to use in kinase activity assays. All complexes were dephosphorylated during purification to remove any initial heterogenous phosphorylation. The SIN1-S69A autophosphorylation site mutant did not affect AKT-mediated activation of mTORC2, suggesting autophosphorylation (and thus ordering of the SIN1 string) is not a critical step in the activation mechanism (**Fig. 3G**). Interestingly, the SIN1-T86A mutant resulted in a slight increase in basal mTORC2 activity (**Fig. 3G**). We hypothesize that the T86A mutation is sufficient to partially dislodge the SIN1 bridge, as the T86A substitution would be unable to form autoinhibitory SIN1-T86/Rictor-E1675 interaction observed in our structure. Despite this increase in basal activity, SIN1-T86A mTORC2 was largely insensitive to further activation by AKT^WT^ (∼2-fold activation) when compared to SIN1 WT or S69A mTORC2 (∼30-fold activation; **Fig. 3G and 3H**), highlighting the importance of SIN1-T86 phosphorylation for full activation of mTORC2. Double mutation of both SIN1 AKT phosphorylation sites (SIN1-T86A/S510A) behaved similarly to T86A alone, showing that the modest 2-fold activation of T86A mutants by AKT^WT^ is not driven by phosphorylation at SIN1-S510.

Furthermore, Phos-tag analysis of the SIN1-T86A/S510A mTORC2 incubated with ATP + AKT^WT^ resulted in SIN1 autophosphorylation only, confirming that T86 and S510 are the only AKT target sites within SIN1 (**fig. S9A**), suggesting the low-level AKT^WT^ activation of SIN1-T86A containing mutants is via a SIN1 phosphorylation-independent mechanism.

### Dynamic structural changes in mTOR active site upon AKT-phosphorylation

To further investigate dynamic structural changes during mTORC2 activation by AKT, we performed hydrogen deuterium exchange coupled to mass spectrometry (HDX-MS) on dephosphorylated, auto-phosphorylated and AKT phosphorylated mTORC2. We observed changes in deuterium incorporation in the SIN1/Rictor interface and the mTOR active site in both autophospho- and phospho-mTORC2 versus dephospho-mTORC2 (**Fig. 4**). Firstly, the SIN1 string-Rictor interface is protected in both auto-and phospho-mTORC2 (**Fig. 4A-C**): Rictor helices α2 and α4 at the interface with the SIN1 string showed a strong protection, supporting our cryo-EM structural results where autophosphorylation of SIN1-S69 leads to the SIN1 string anchoring on Rictor helices α2/α4/α7. Additional, light protection is observed for Rictor helices α1, α10 and α12 in the vicinity of the SIN1 NTD. In SIN1, a strong protection is observed at the C-terminus of the α1-α2 helical hairpin (residues 33-38), in both autophospho- and phospho-mTORC2 (**Fig. 4D and E**), where our cryo-EM maps show a few additional ordered SIN1 residues. Additionally, a light deprotection is observed in the Rictor CTD-α3 helix in phospho-mTORC2 only, suggesting increased flexibility of this region (**Fig. 4C**), consistent with the observed disordering of the SIN1 bridge upon AKT-phosphorylation and loss of interaction of SIN1-R89 in the bridge with E1633 in Rictor CTD-α3 within an acidic patch of Rictor.

**Fig. 4.**
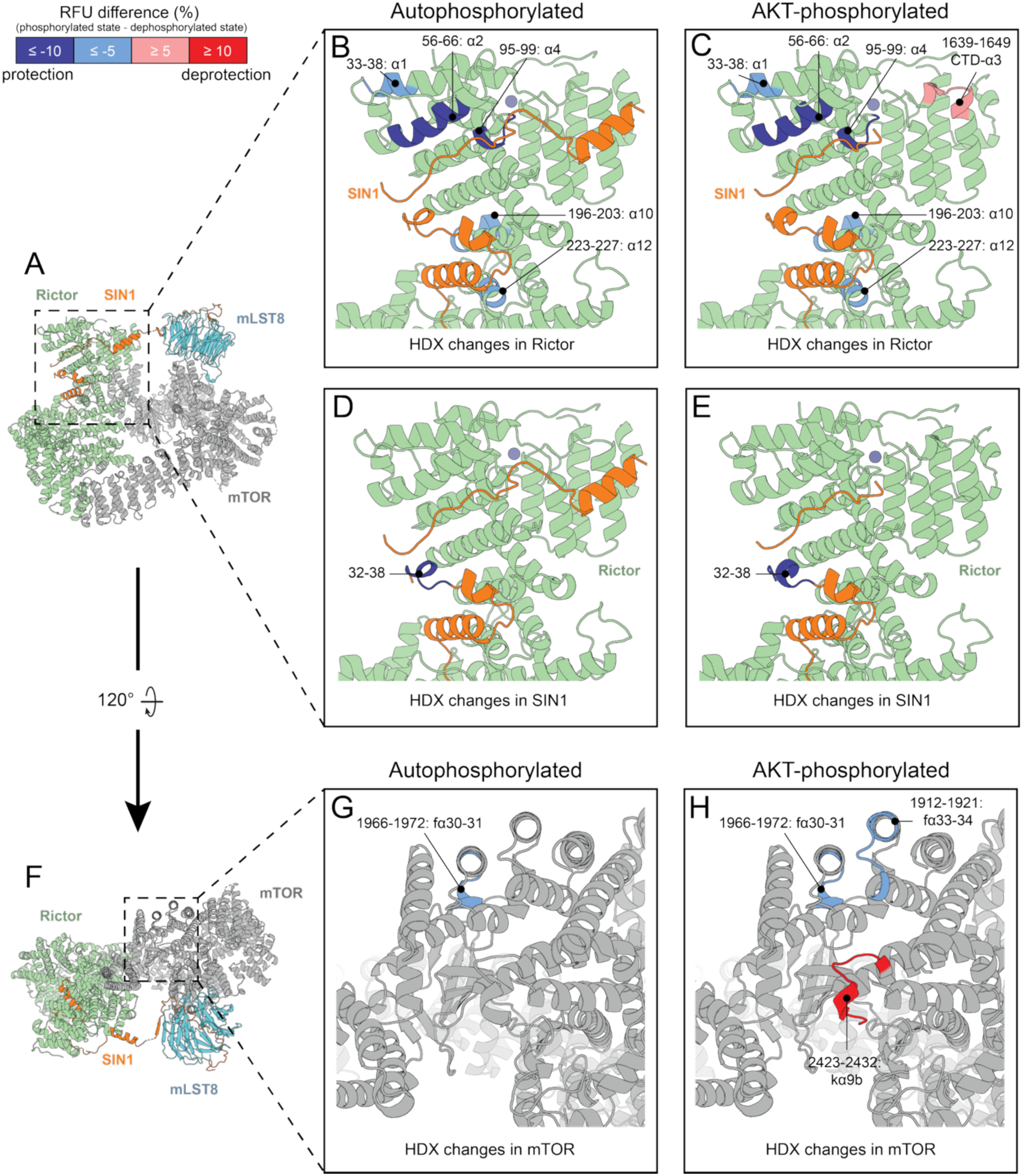
Phosphorylation by AKT reorganizes the mTORC2 active site. HDX-MS characterization of dephosphorylated, auto-phosphorylated and AKT-phosphorylated mTORC2. Validated relative fractional uptake (RFU) differences (autophosphorylated vs dephosphorylated and AKT-phosphorylated vs dephosphorylated) are illustrated on the AKT-phosphorylated and autophosphorylated mTORC2 structures. (A-E) HDX-MS changes at the SIN1/Rictor interface. Rictor and SIN1 peptides showing statistically validated changes in HDX upon autophosphorylation (Rictor peptides in panel B and SIN1 peptides in panel D) and AKT-phosphorylation (Rictor peptides in panel C and SIN1 peptides in panel E) are shown. (F-H) HDX-MS changes in the mTOR ATP binding site. mTOR peptides showing statistically validated changes upon autophosphorylation and AKT-phosphorylation (G and H respectively) are shown. HDX changes are colored as follows: blue for protection and red for deprotection.

Secondly, a robust impact is observed in the mTOR active site upon AKT phosphorylation of mTORC2 (**Fig. 4F-H**). A strong deprotection is observed in helix kα9b only in mTORC2 phosphorylated by AKT (**Fig. 4H**). kα9b is localized in the catalytic cleft and overlaps with the PIKK regulatory domain (PRD), deletion of which increases mTOR activity (*40–43*). These data suggest that AKT phosphorylation may drive reorganization of the kα9b/PRD region within the active site as an additional driver of mTORC2 activation. Some light protection is observed in mTOR helices fα30-31 upon auto-phosphorylation (**Fig. 4G**) and in helices fα30-31-33-34 upon AKT-phosphorylation (**Fig. 4H**). Interestingly, these helices are part of the mTOR dimer interface where a smaller inner diameter of the space between the two protomers has been observed in our corresponding cryo-EM maps.

### A phosphorylation driven AKT-mTORC2 positive feedback loop

As mTORC2 is a critical activator of AKT via phosphorylation of the AKT C-terminal tail, and we have in turn shown AKT to be a potent activator of mTORC2, we next sought to investigate this putative mTORC2-AKT positive feedback loop in our *in vitro* reconstituted system. To monitor simultaneous activation of both mTORC2 and AKT in the same experiment, we first combined dephosphorylated mTORC2 and dephosphorylated AKT^WT^ at high (1 μM) concentrations, in the presence of ATP, to allow reciprocal phosphorylation/activation to occur. The status of phosphorylation of mTORC2 (measured by SIN1-pT86) and AKT^WT^ (measured by AKT-pS473) after this prephosphorylation reaction was examined by western blotting. In tandem, the pretreated enzymes were seeded at low concentration (30 nM) into separate kinase assays that measured either mTORC2 activity (using kinase-dead AKT^D274A^ as a substrate) or AKT kinase activity (using GST-tagged SIN1 PH as a substrate, based on our observation that SIN1-S510 is an AKT target site) (**Fig. 2E and 3F**), allowing correlation of mTORC2 and AKT phosphorylation state with their respective kinase activities. When inputs for the pre-phosphorylation reaction were both mTORC2 and AKT in the dephosphorylated state, even after the prephosphorylation reaction, both enzymes had very low activity, as judged by the kinase assays which show lack of phosphorylation of either mTORC2 substrate (AKT^D274A^, lanes 1’-4’) or AKT substrate (GST-PH, lanes 1”-4”) (**Fig. 5A**). This low kinase activity correlated with only limited amounts of phosphorylated mTORC2 (SIN1-pT86, lanes 1-4) and incomplete phosphorylation of AKT (AKT-pS473). We hypothesized this low activity was due to the requirement for phosphorylation of AKT-T308 within its activation loop by PDK1 (*44–46*). Indeed, treatment of dephospho-AKT with PDK1 to generate AKT-pT308 was sufficient to stimulate AKT activity, even in the absence of mTORC2 treatment (GST-PH, **Fig. 5A**, lane 7”). When AKT-pT308 was included in prephosphorylation reactions with mTORC2, the mTORC2 activity was dramatically increased (pAKT^D274A^, lane 8’) and a further enhancement of AKT activity beyond AKT-pT308 alone was observed (pGST-PH, **Fig. 5A**, lane 8”). This enhanced activation of AKT coincided with phosphorylation of mTORC2 (SIN1-pT86, lane 8). Therefore, initial stimulation of AKT by PDK1 phosphorylation at pT308 is a prerequisite for positive feedback via an AKT-pS473/mTORC2 SIN1-pT86 positive feedback loop.

**Fig. 5.**
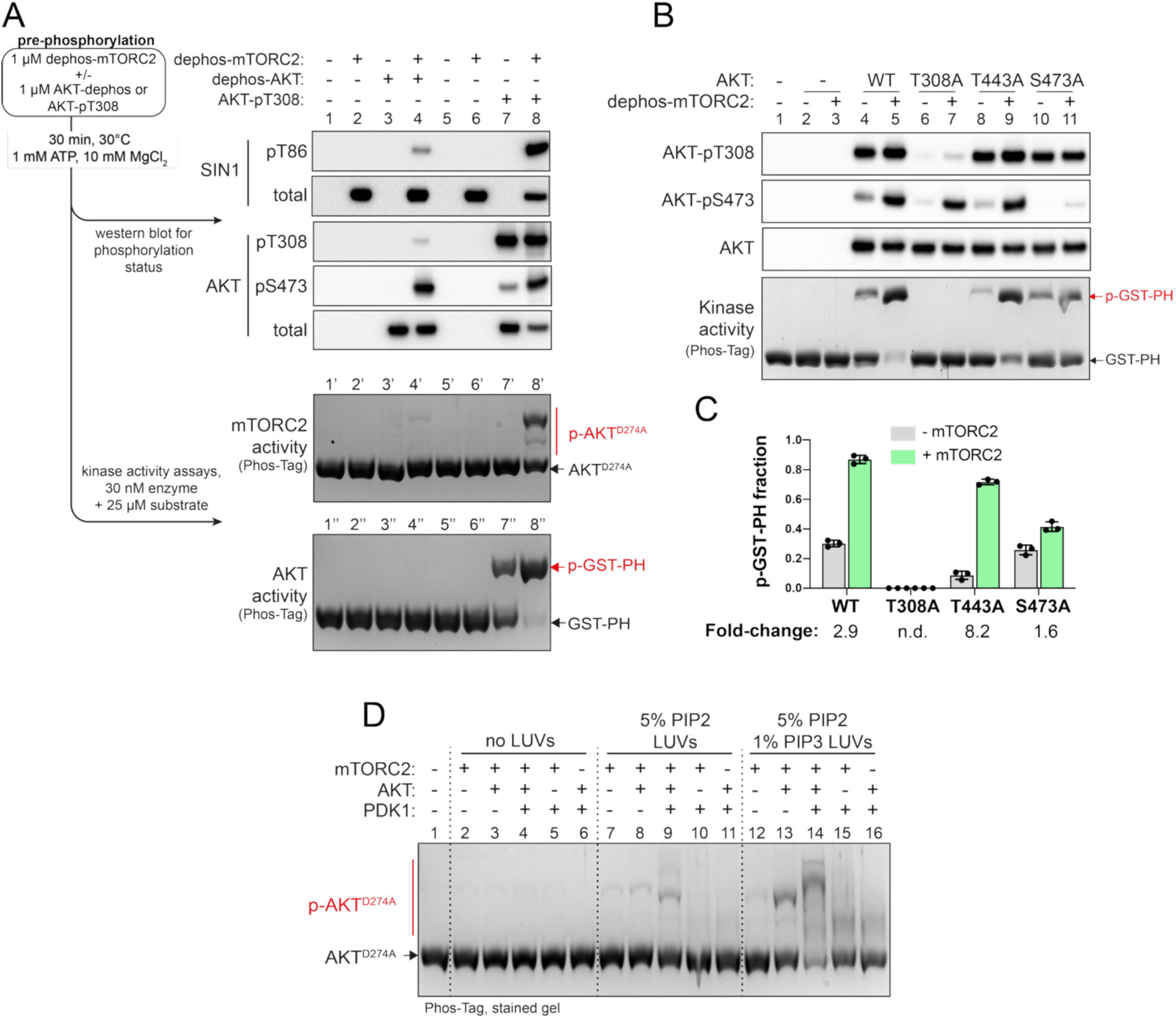
A membrane localized mTORC2-AKT positive feedback loop. (A) Simultaneous activation of mTORC2 and AKT by positive feedback. 1 µM of the indicated combinations of dephosphorylated (dephos)-mTORC2 with either dephos-AKT or AKT-pT308 were incubated with 1 mM ATP, 10 mM MgCl2 for 30 min at 30°C. Reaction mixes were then analyzed by western blotting for their phosphorylation status (top) or seeded into kinase assays at 30 nM using 25 µM of the indicated substrates for 10 min at 30°C to measure to corresponding enzymatic activity (bottom). (B) Kinase activity of AKT mutants after mTORC2 phosphorylation. 1 µM of the indicated AKT mutants were incubated with 1 µM dephos-mTORC2 for 30 min at 30°C and resulting AKT phosphorylation state and kinase activity determined as described for (A). (C) Quantification of the fraction of GST-SIN1-PH phosphorylated by AKT mutants in (B). Data represents the mean of n = 3 independent experiments ± SD. The mean fold-change upon treatment with mTORC2 (+mTORC2/-mTORC2) is indicated below for each mutant. (D) PIP-containing membranes are required for efficient activation of AKT/mTORC2 signaling. The indicated combinations of dephosphorylated enzymes (all 30 nM) and liposomes (0.4 mg/mL) were used in mTORC2 activity assays with 25 µM AKT^D274A^ substrate for 10 min at 30°C.

Recently, it has been proposed that activation of both PKC and AKT by mTORC2 is stimulated via phosphorylation of a ‘TOR interaction motif’ (TIM; T443 in AKT1), which activates AKT and allows hydrophobic motif (HM) phosphorylation at S473 by AKT autophosphorylation (*10*). To further investigate the contribution of specific AKT phosphorylation sites to the AKT-mTORC2 positive feedback loop, we generated alanine mutants at key phosphorylation sites within AKT for use in activity assays: the PDK1 activation loop site T308A, the putative TIM site T443A, and the HM site S473A. AKT^WT^ and mutant proteins were dephosphorylated, then incubated with PDK1 to enable the first activation step via T308 phosphorylation; the TIM and HM mutants were phosphorylated on AKT pT308 to a similar level as AKT^WT^ (**Fig. 5B**). The kinase activity of AKT (using GST-PH as a substrate) was then examined with or without prephosphorylation by mTORC2, along with corresponding phosphorylation at AKT pS473 (**Fig. 5B and C**). AKT^WT^ had basal kinase activity (due to phosphorylated pT308) that could be significantly stimulated by mTORC2 treatment, coinciding with phosphorylation of AKT-pS473 (lane 5 vs lane 4) (**Fig. 5B and C**). The AKT-T443A TIM mutant had lower basal activity relative to AKT^WT^ (lane 8 vs lane 4) but could still be efficiently activated by incubation with mTORC2 (lane 8 vs lane 9), again coinciding with generation of AKT-pS473 (**Fig. 5B and C**), suggesting that T443 phosphorylation is not essential for AKT activation by mTORC2. Critically, the HM AKT-S473A mutant had comparable basal activity to AKT^WT^ (**Fig. 5B**, lane 10 vs lane 4) but could not be fully activated by incubation with mTORC2 (lane 11 vs 10) (**Fig. 5B and C**). As expected, AKT-T308A had no detectable basal activity (i.e., it could not be activated by PDK1) and could not be activated by mTORC2 (lane 7 vs lane 6). We did observe detectable pS473 phosphorylation of the inactive AKT-T308A mutant upon incubation with mTORC2 (**Fig. 5B**), again suggesting that phosphorylation of the hydrophobic motif is not a product of AKT autophosphorylation (AKT-pS473, **Fig. 5B**, lane 7). Similarly, we found that mTORC2 could phosphorylate the HM S473 within kinase dead AKT^D274A^ (**fig. S11A and S11B**), further showing AKT activity (and thus potential autophosphorylation) is not required for phosphorylation of the hydrophobic motif. Together, these results show that mTORC2 is required for full activation of AKT, driven via phosphorylation of the AKT hydrophobic motif S473 by mTORC2.

In addition to activation by phosphorylation, both PDK1 and AKT1 undergo autoinhibition via their PH domains, which is relieved by PIP3 binding (*47–50*). To test the importance of membrane localization in this mTORC2-AKT positive feedback loop, we reconstituted the full cellular system *in vitro* using PDK1, AKT and mTORC2 on LUVs. For these experiments all proteins were used at 30 nM in kinase assays, which were judged to be near physiological concentration (**Table S4**). At 30 nM concentration, in the absence of membrane, no mTORC2 activity could be observed in kinase assays even in the presence of AKT^WT^ and PDK1 (**Fig. 5D**, lanes 1-6). mTORC2 activity was moderately enhanced by 5% PIP2 liposomes in line with our results showing membrane activation of mTORC2 (**Fig. 5D**, lane 2 vs 7 and **Fig. 1B**). This was no further activated by incubation with AKT^WT^ alone but was further stimulated by AKT^WT^ + PDK1 (**Fig. 5D**, lane 8 vs 9). The inclusion of 1% PIP3 in 5% PIP2 liposomes, simulating PIP3 generation by PI3K, lead to stronger mTORC2 activity when using AKT^WT^ and mTORC2 alone (**Fig. 5D**, lane 13), likely due to disinhibition of AKT by PH domain binding to PIP3 allowing mTORC2 phosphorylation, as well as co-localization of mTORC2 and AKT on the membrane by PIP2 and PIP3, respectively. Maximum activation of mTORC2 was achieved on PIP2/PIP3-containing liposomes by inclusion of PDK1 together with AKT^WT^ (**Fig. 5D**, lane 14), in agreement with a combined effect of all 3 components now being co-localized on PIP2/PIP3-containing membranes, disinhibition of AKT1 and PDK1 PH domains by PIP3 and mTORC2 activation by PIP2. In line with our observation that mTORC2ΔPH retains binding to PIP2 membranes (**fig. S2**), we found the SIN1-PH was dispensable for activation of mTORC2, with mTORC2ΔPH being efficiently activated by AKT and PDK1 in a PIP2/PIP3 dependent manner similar to WT mTORC2 (**fig. S12A and S12B**). These experiments at near physiological concentrations highlight that membrane localization of mTORC2, along with the signaling partners AKT and PDK1, is key for efficient activation of the AKT-mTORC2 positive feedback loop.

In summary, these results provide a clear mechanism for the activation of growth-factor mediated mTORC2 activity, driven via an AKT-mTORC2 positive feedback loop:

1) mTORC2 is resident at cellular membranes via mTORC2-PIP2 interactions. 2) Growth factors induce PI3K-mediated generation of PIP3 in the inner-leaflet. 3) PIP3 recruits PDK1 and AKT1 to the membrane. 4) Activated PDK1 phosphorylates the AKT1-pT308 activation loop and partially activates it. 5) Partially active AKT1 phosphorylates and activates mTORC2 via SIN1-pT86. 6) Activated mTORC2 phosphorylates and fully activates AKT1 via pS473. 7) AKT1 is fully activated and can activate further mTORC2 molecules to enhance positive feedback and/or phosphorylate downstream substrates.

## Discussion

In this study we have investigated the activation mechanisms of mTORC2 in a reconstituted system of PI3K signaling, consisting of PIP-containing membranes, mTORC2, AKT and PDK1. We find that mTORC2 alone robustly binds to PIP2-containing membranes, even in the absence of the SIN1 PH domain, and our cryo-ET reconstruction of mTORC2 alone on PIP2 containing LUVs revealed the orientation of mTORC2 on the membrane, highlighting the mTOR N-HEAT domain as the major membrane interface within the complex. The overall orientation of mTORC2 in relation to the membrane is similar to that observed for the membrane-bound mTORC1-Rag-Ragulator complex (*38*). In mTORC2, the membrane proximal L1-L6 loops in the N-terminal segment of the mTOR N-HEAT (**Fig. 1E and fig. S4**) are the primary contact sites with membrane, with basic residues in these loops located near the membrane. Although mTORC1 faces the membranes in a similar way as mTORC2, direct interaction of the mTOR subunit with membranes within mTORC1 is limited to only loop L6 (mTOR residues 470-475, **Fig. 1E and fig. S4 and S13**).

We further observed the Rictor subunit in contact with the membrane, including a basic patch that is part of the Rictor ‘A-site’ (basic residues K541, R575, R576, and R572) that was previously observed to bind ATP (*21*). It is appealing to propose that on the membranes, PIP2 might be an alternative negatively charged ligand for the ‘A-site’, since the Rictor A-site is oriented toward the membrane in a manner reminiscent of a membrane-facing hydrophobic loop within the Raptor subunit of mTORC1 (Raptor F1296/M1297), which plays a role in membrane binding of the mTORC1-Rag-Ragulator-RHEB complex (*38*) (**fig. S13**). Mutations which abolish nucleotide binding in the Rictor A-site *in vitro* had no effect on mTORC2 signaling in cells (*21*), perhaps due to retention of the major membrane contact site of the mTOR N-HEAT described here. The orientation of mTORC2 that we observe in our cryo-ET reconstruction is inverted with respect to the orientation that was recently proposed for a complex of mTORC2 bound to AKT, based on crosslinking data and computational modelling (*28*). Majority of mTORC2 substrates are membrane interacting proteins on their own, so it is plausible that they can modify arrangement of mTORC2 on membranes, and/or the extent of mTORC2 activation by membranes. Future cryo-EM studies with mTORC2 on the membranes in the presence of the membrane-interacting substrates will allow comparison with apo mTORC2 on membranes described here.

We found no clear role for the SIN1-PH domain in membrane binding or activation of mTORC2, with mTORC2ΔPH capable of membrane interaction, as well as being fully activated by AKT phosphorylation in a reconstituted system. It has previously been proposed that the SIN1-PH forms an autoinhibitory interaction with the mTOR kinase domain within mTORC2, which is relieved by binding to PIP3. While we confirm SIN1-PH can bind both PIP2 and PIP3, we observed no density for PH domain interaction with mTORC2 (cis-) or membranes (trans-) in either SPA or tomography reconstructions. Activity of mTORC2 within a cell has been shown to localize to many sub-cellular compartments including the plasma membrane, mitochondria, endosomes and the ER (*12, 15*). Therefore, the cellular role of the SIN1-PH could be in trafficking of mTORC2 to specific sub-cellular membranes, to be localized alongside activators such as PDK1/AKT, a nuance which is not required in our *in vitro* reconstituted system. Indeed, determining how mTORC2 activity is compartmentalized between specific sub-cellular membranes remains a key open question. In contrast to the stepwise recruitment of mTORC1 to lysosomal membrane - which is mediated by membrane-anchored interacting partners Rag/Ragulator and Rheb, followed by Raptor-membrane interactions and finally mTOR-membrane interactions (*38*) - the intrinsic binding activity of mTORC2 to membranes may be sufficient to drive its membrane localization in cells.

Our biochemical and structural observations highlight a critical mechanistic role for AKT phosphorylation of T86 within the SIN1 bridge in activation of mTORC2, with phosphorylation driving an order-disorder transition. It has been previously reported that SIN1 T86 and T398 can be phosphorylated by S6K, leading to inhibition of mTORC2 by dissociation of SIN1 from the complex (*33*), although it has also been argued that AKT is the physiological T86 kinase (*35*). We did not observe SIN1-T398 phosphorylation in our study, which is localized to the β3 strand of the SIN1-PH domain, and this may indeed be an S6K specific site. In line with our observations, in previous mTORC2 structures it has been noted that T86 occupies a negatively charged pocket in Rictor, and it was proposed that phosphorylation would repulse T86 to disrupt the complex (*21*). We now show that SIN1 T86 phosphorylation leads to localized flexibility within the SIN1-string/bridge-Rictor interaction region and activation of mTORC2, rather than SIN1 dissociation and mTORC2 inhibition.

The SIN1 string region has been implicated previously in mTORC2 substrate recognition, being critical for recruitment of SGK1 but not AKT (*39*). Whilst we found a role for SIN1-pS69 autophosphorylation in anchoring of the SIN1-string to the Rictor ARM domain, we saw no effect on mTORC2 stability or activity in an autophosphorylation-impaired SIN1-S69A mutant, although we focused on AKT as a substrate and therefore may be missing SGK1 specific regulation. Indeed, determining whether mTORC2 activation by phosphorylation is specific to signaling via AKT only or whether it can be performed by other AGC kinase mTORC2 substrates is an important area for future work. Our *in vitro* activity assays and structures highlight a key role for T86 phosphorylation, which we show is necessary for mTORC2 activation. These observations highlight that the phosphorylation state is a key consideration in interpretation of mTORC2 structure and function in future studies.

We further observed dynamic changes in the mTORC2 active site by HDX-MS, showing increased flexibility within the mTOR kα9b in response to AKT phosphorylation. In the PIKK family member ATM, kα9b within the PRD blocks substrate binding in the active site, which must then be removed during activation (*51*). In mTORC1, the PRD is also proposed to restrict active site access, with kα9b partial deletion leading to mTORC1 hyperactivity (*41, 43*). Molecular dynamics simulations of mTOR with and without deletion of the PRD suggest this region bridges the catalytic cleft and forms interactions with the FRB, limiting conformation flexibility and substrate access (*52*). Strikingly, the SIN1 bridge also spans the mTORC2 catalytic cleft and interacts with the Rictor CTD, which sits atop the mTOR FRB. Therefore, it could be hypothesized that AKT phosphorylation drives both dislodging of the SIN1 bridge from the Rictor CTD and increased mobility in the mTOR PRD, leading to increased conformational flexibility, substrate access and thus kinase activation.

While we show mTORC2 *k*cat was activated 2-fold by membrane binding, this is modest compared to the >20-fold activation of mTORC2 driven by AKT phosphorylation. Despite this, we show a critical requirement for mTORC2 membrane localization in efficient signaling. At physiological concentrations, even in the presence of the AKT activating kinase PDK1, the AKT-mTORC2 positive feedback loop could not be activated in the absence of PIP2/PIP3 membranes. Both AKT and PDK1 require disinhibition of their PH domains for full activation (*53*), with PDK1 also undergoing membrane-localized autophosphorylation during activation (*49*). Therefore, it seems likely that efficient activation of PI3K signaling is driven by a combination of factors: localized increases in substrate concentration by co-localization and reduction of dimensionality to a planar membrane, combined with activation of mTORC2 by PIP2 binding, activation of PDK1 and AKT by PIP3, and phosphorylation-dependent mutual activation of mTORC2 and AKT.

## Acknowledgments

We thank Anna Yeates, Grigory Sharov, Giuseppe Cannone of the MRC-LMB EM facility for help with cryo-EM data collection, Jake Grimmett, Toby Darling and Ivan Clayson for assistance and advice with scientific computing, Stephen McLaughlin for help with biophysical instrumentation, and Tomos Morgan, Sew-Yeu Peak-Chew, Farida Begum and Catarina Franco from the BioMass Facility for help with MS experiments.

## Author contributions

Conceptualization: IMH, OP, RLW

Methodology: IMH, MB, BA, OP, MA, RLW

Investigation: IMH, OP, MB

Visualization: IMH, MB

Funding acquisition: RLW

Project administration: RLW Supervision: RLW

Writing – original draft: IMH

Writing – review & editing: IMH, OP, RLW, MB, MA

### Data and materials availability

All materials are available upon request from the corresponding author. Atomic models and cryo-EM density maps were submitted to the PDB and EMDB as follows. Dephosphorylated-mTORC2: PDB ID 9TDS and EMD-55803 (composite dimer), PDB ID 9TDT and EMD-55804 (high-resolution focused protomer); autophosphorylated-mTORC2: PDB ID 9T92 and EMD-55714 (composite dimer), PDB ID 9T93 and EMD-55715 (high-resolution focused protomer); AKT-phosphorylated-mTORC2: PDB ID 9T7J and EMD-55637 (composite dimer), PDB ID 9T94 and EMD-55716 (high-resolution protomer). The raw consensus cryo-EM dimer maps can be accessed as related entries to the composite EMDB submissions. The atomic model and cryo-ET subtomogram averaged map for mTORC2 on PIP2 containing membrane were deposited in the PDB and EMDB under accession codes 9TPW and EMD-56117, respectively.

## Supplementary Materials

Materials and Methods

Figs. S1 to S13

Tables S1 to S4

References (54-73)

