## Supplementary Information for "Structural basis for a phosphoinositide-driven mTORC2-AKT positive feedback loop"

### Supplementary Materials

### Materials and Methods

#### Plasmids and constructs

Human mTORC2 subunits were cloned individually into a mammalian expression vector as described previously (*54*). The mTOR subunit, with an N-terminal tandem Strep-tag II followed by a TEV cleavage site, was cloned into a pCAG vector. The mLST8 subunit without a tag was cloned into a pCAG vector. Human SIN1.1, PCR-amplified from Addgene 73388 plasmid (gift from Jie Chen and Taekjip Ha (*55*)), was cloned without a tag into pcDNA4TO. Next, the promoter-gene (SIN1.1)-terminator cassette was PCR amplified from this plasmid and cloned into a pCAG vector already containing mLST8 gene, enabling co-expression of SIN1 and mLST8 from a single plasmid. Human RICTOR, PCR-amplified from IMAGE:9021161 clone, was cloned with a 3XFlag-tag or without a tag in the pCAG vector. The SIN1 phospho-site mutants were made by overlapping PCR mutagenesis and cloned into a pCAG vector already containing mLST8. The SIN1-PH domain (residues 370-522) gene fragment was synthesized (IDT gBlock) and cloned into a pOPTG vector containing an N-terminal GST tag and TEV cleavage site.

Wild-type AKT1 with an N-terminal $His_{10}$-StrepII-(tev) tag, cloned in the pFastBacDual baculovirus expression vector, was a gift from Thomas Leonard, Addgene plasmid 86561 (*55*). Human AKT1_D274A was made by overlapping PCR mutagenesis using wild-type AKT1 as a template and cloned with an N-terminal tandem StrepII-(tev) tag into pAceBac1. All AKT1 phospho-site mutations were made using overlapping PCR mutagenesis and were cloned with tandem StrepII-(tev) tag in the pAceBac1.

#### Protein expression

Expression of mTORC2 complexes was performed by transient transfection of Expi293F cells, cultured at 37°C, 8% CO2 with orbital shaking at 125 rpm. Cells were co-transfected with a total of 1.1 mg of DNA/L cells at a density 2.5 x $10^6$ cells/mL using polyethylenimine (PEI, 3 mg PEI/L cells). Cells were harvested by centrifugation 68 h post-transfection and snap-frozen in $LN_2$. All recombinant AKT constructs were expressed for 49 h in Sf9 cells.

GST-SIN1_PH was expressed in *Escherichia coli* OverExpress C41(DE3) cells. Cells transformed with the expression plasmid were cultured in 2X TY medium at 37°C until $OD_{600}$ = 0.6 and protein expression induced with 0.5 mM isopropyl-β-D-1-thio-galactopyranoside for 18 h at 20°C. Cell pellets were harvested by centrifugation and snap-frozen in $LN_2$.

**Protein purification**

All mTORC2 constructs were purified using the following general protocol. Key differences for producing differentially phosphorylated mTORC2 states are noted at the appropriate step. All protein lysis and chromatography steps were performed at 4°C. Cell pellet from 2 L of mTORC2-Expi293 cells was resuspended in 200 mL of ice-cold lysis buffer (50 mM bicine, pH 8.5, 300 mM NaCl, 2 mM $MgCl_2$, 0.5 mM TCEP) supplemented with 1X EDTA-free protease inhibitors, 25 U/mL Pierce universal nuclease and 0.1 mM PEFA. Cells were lysed by sonication at 40% amplitude for 2 min at 4°C and lysate was clarified by centrifugation for 30 min at 35,000 RPM, 4°C in a Ti45 rotor. The clarified cell lysate was loaded onto 2 tandem StrepTrap HP 5 mL columns, then washed with 6 CV lysis buffer followed by 6 CV wash buffer (50 mM BICINE, pH 8.5, 150 mM NaCl, 2 mM EDTA, 0.5 mM TCEP) and protein eluted with wash buffer supplemented with 10 mM desthiobiotin. For untreated mTORC2, protein was immediately purified by anion exchange chromatography (AEX). Protein was diluted to <100 mM NaCl with Q buffer A (50 mM bicine, pH 8.5, 50 mM NaCl, 0.5 mM TCEP) and loaded onto a HiTrap Q 5 mL column equilibrated in Q buffer A. Protein was eluted by a 0-100% linear gradient of Q buffer B (Q buffer A with 1 M NaCl). Fractions containing mTORC2 were pooled and concentrated to ~7 mg/mL in a 100 kDa cut-off centrifugal filter and snap-frozen in $LN_2$.

For dephosphorylated mTORC2, after AEX purification, protein was concentrated to ~2 mL, supplemented with 1 mM $MnCl_2$ and incubated with lambda protein phosphatase (λ-PP, ~2 µg of phosphatase per mg of mTORC2) overnight at 4°C. To remove λ-PP, the dephosphorylated protein was then further purified by size-exclusion chromatography (SEC) on a Superose6 16/600 column equilibrated in SEC buffer (50 mM bicine, pH 8.5, 250 mM NaCl, 1 mM TCEP) then concentrated and stored as for untreated mTORC2.

For AKT phosphorylated mTORC2, dephosphorylation was performed on-column after protein binding to the StrepTrap column by injecting 1 CV λ-PP (50 µg total) in wash buffer supplemented with 1 mM $MnCl_2$. After overnight incubation at 4°C, the column was washed with 2 CV wash buffer to remove λ-PP before elution with desthiobiotin as described above. Protein was purified by AEX as above and concentrated to ~2 mL. This dephosphorylated mTORC2 was then phosphorylated *in vitro* with recombinant pT308-AKT$^{WT}$ (purified as described below, pre-phosphorylated with PDK1 *in vitro*) at a 1:1 mTORC2:AKT ratio supplemented with 1 mM ATP, 10 mM $MgCl_2$, for 1 h at 30°C. Finally, mTORC2 was purified by SEC, concentrated and stored as described for dephospho-mTORC2.

For purification of AKT$^{D274A}$, cell pellet from 2 L of AKT$^{D274A}$-Sf9 cells was resuspended in 200 mL lysis buffer (50 mM Tris-HCl, pH 8.0, 300 mM NaCl, 0.5 mM TCEP) supplemented with 1X EDTA-free protease inhibitors, 25 U/mL Pierce universal nuclease and 0.1 mM PEFA. Cells were lysed by sonication, lysates clarified and protein purified by StrepTrap affinity chromatography as described for mTORC2 Expi293 purification. For AKT, the wash buffer consisted of 50 mM HEPES, pH 7.8, 150 mM NaCl, 0.5 mM TCEP and elution was carried out in wash buffer supplemented with 10 mM desthiobiotin. The pooled AKT fractions were supplemented with 1 mM TCEP, 1 mM $MnCl_2$ for simultaneous overnight TEV protease cleavage/λ-PP dephosphorylation, using 1:30 (TEV:AKT) by mass and 1 µg λ-PP per 1 mg AKT. The cleaved, dephosphorylated AKT was further purified by AEX on a HiTrap Q 5 mL column as described for mTORC2, using Q buffers consisting of 50 mM HEPES, pH 7.8, 25 mM (A buffer) or 1 M (B buffer) NaCl, 0.5 mM TCEP. Fractions containing AKT$^{D274A}$ were pooled and concentrated to ~30 mg/mL in a 30 kDa cut-off centrifugal filter and snap-frozen in $LN_2$.

Purification of AKT$^{WT}$ (and 'active' mutants) was performed as for AKT$^{D274A}$ with the following changes. The lysis buffer was supplemented 50 mM NaF and no dephosphorylation was performed during TEV cleavage. Wash and Q buffers were as for AKT$^{D274A}$ except for usage of Tris pH 8.0 in all steps. Dephosphorylated AKT$^{WT}$ was produced by *in vitro* λ-PP dephosphorylation of defrosted AKT$^{WT}$ followed by SEC purification. AKT$^{WT}$-pT308 was produced by *in vitro* phosphorylation of AKT$^{WT}$ with PDK1 and isolated by AEX on a MonoQ 5/50 GL column.

For purification of GST-SIN1_PH, cell pellet from 2 L *E. coli* culture expressing GST-SIN1_PH was resuspended in 50 mL lysis buffer (50 mM Tris-HCl, pH 8.0, 500 mM NaCl, 5% [v/v] glycerol, 0.5 mM TCEP) supplemented with 1X EDTA-free protease inhibitor cocktail, 0.01 mg/mL DNase, 1 mg/mL lysozyme and 0.1 mM PEFA. Cells were lysed by sonication at 75% amplitude for 4 min at 4°C and clarified by centrifugation for 30 min at 35,000 RPM, 4°C in a Ti45 rotor. The clarified lysate was incubated with washed glutathione sepharose 4B beads for 1 h at 4°C. Beads were packed into a gravity column, washed with 10 bed-volumes of lysis buffer followed by 5 bed-volumes of wash buffer (50 mM

Tris-HCl, pH 7.4, 100 mM NaCl, 5% [v/v] glycerol, 0.5 mM TCEP). Protein was then eluted in fractions of wash buffer supplemented with 50 mM reduced glutathione. Protein containing fractions were pooled, diluted to <50 mM NaCl in S buffer A (50 mM Tris-HCl, pH 7.4, 25 mM NaCl, 0.5 mM TCEP) and loaded onto a HiTrap SP 5 mL cation exchange column equilibrated in S buffer A. Bound protein was eluted by a 0-100% linear gradient against S buffer B (S buffer A with 1 M NaCl). Protein containing fractions were pooled, concentrated to 2 mL in a 30 kDa cut-off centrifugal filter and purified further by SEC on a s200 16/600 column equilibrated in SEC buffer (25 mM Tris-HCl, pH 7.4, 150 mM NaCl, 0.5 mM TCEP). Protein containing fractions were pooled and concentrated to ~20 mg/mL and snap-frozen in $LN_2$.

**LUV preparation**

Lipid mixes used for LUVs are described in **Table S1**. PIP2 and PIP3 lipids were resuspended at 1 mg/mL in a mixture of chloroform, methanol and water (20:9:1 by volume), all other lipids were purchased as chloroform stocks. Lipids were mixed in glass vials and dried to a lipid film by rotating the vial under a stream of nitrogen gas. Lipid films were completely dried in a vacuum desiccator for 1 h. Lipids were then resuspended in lipid buffer (25 mM HEPES, pH 7.5, 150 mM KCl, 1 mM TCEP) by vortex for 3 min followed by incubation in a sonicating water bath for 5 min. The lipid mixes were then transferred to plastic 1.5 mL tubes and sonicated to clarity in a non-contact sonicator (VialTweeter, Hielscher; ~6 x 10 s pulses for 5 mg/mL lipid stocks). Samples were then freeze-thawed for 10 cycles between $LN_2$ and a 45°C water bath. Finally, samples were passed 15 times through a 100 nm syringe filter (Whatman Anotop) and either used immediately or snap-frozen in $LN_2$.

**LUV flotation assays**

For mTORC2 flotation assays, 2 µM mTORC2 was mixed with 1.4 mg/mL LUVs in a total volume of 20 µL made up in flotation buffer (25 mM HEPES, pH 7.5, 100 mM NaCl, 0.5 mM $MgCl_2$, 0.5 mM TCEP) and incubated on ice for 30 min. The 20 µL samples were then mixed with 60% sucrose (w/v, made up in flotation buffer) to a 35% final sucrose concentration. The sample (40 µL) was placed at the bottom of a 0.2 mL thick-wall polycarbonate tube (Beckmann Coulter), followed by careful pipetting on of the following sucrose gradient on top: 52 µL 30% sucrose, 52 µL 25% sucrose, 16 µL flotation buffer. The remaining mTORC2/LUV mixture was retained as the input sample. Sucrose gradients were then centrifuged for 1.5 h at 258,000 x *g* in a TLS-55 rotor (Beckmann Coulter). After centrifugation, 6 x 26 μL fractions were carefully collected from the top of the gradient for analysis by SDS-PAGE on a NuPAGE 4-12% bis-tris gel using MES buffer, followed by Coomassie staining.

**Kinase assays**

All assays and dilutions were performed in kinase buffer consisting of 25 mM HEPES, pH 7.5, 100 mM NaCl, 5% (v/v) glycerol, 0.5 mM TCEP. Due to the diverse nature of mTORC2 activation experiments with AKT, specific experimental details used for phosphorylation steps and activity assays have been outlined in either the figure or figure legend for each specific experiment. Routinely, kinase assays and pre-treatment phosphorylation steps were performed using 1 mM ATP and 10 mM $MgCl_2$. For assays containing liposomes, the $MgCl_2$ concentration was lowered to 2 mM. Assays were terminated at the indicated timepoints by addition of 4X LDS sample buffer containing 8 mM $ZnCl_2$.

For kinetic experiments, serial dilutions of $AKT^{D274A}$ or $AKT^{D274A}$-ΔPH substrate (10-150 μM) were performed in kinase buffer, followed by incubation with mTORC2 enzyme (0.3 μM for untreated and 30 nM for phospho-mTORC2) for 5 min on ice. For kinetics in the presence of liposomes, 1 mg/ml LUVs were also added during this incubation step. Samples were then equilibrated to 30°C and reactions initiated by the addition of 1 mM ATP and 10 mM $MgCl_2$ (2 mM for liposome kinetics). For each AKT concentration, 1 μg total of AKT was analyzed by Phos-Tag gel. To determine levels of substrate phosphorylation, Coomassie stained Phos-Tag gels were imaged using a Bio-Rad ChemiDoc MP imaging system. Levels of non- and phosphorylated AKT bands were quantified by densitometry using Image Lab software (Bio-Rad) and the total phosphorylated fraction was calculated as (phosphorylated)/(non-phosphorylated + phosphorylated) signals. Kinetic parameters $k_{cat}$ and $K_m$ were calculated using the Michaelis-Menten equation from a non-linear regression fit of the data in GraphPad Prism 10.

**Phos-Tag SDS-PAGE**

Analysis of protein phosphorylation by PhosTag gel was performed on either pre-cast SuperSep 50 μM Phos-tag 7.5% acrylamide gels (FujiFilm Wako) or homemade 7.5% bis-tris gels containing 50 μM Phos-bind reagent (APExBIO), 100 μM $ZnCl_2$. Electrophoresis was performed in 1X MOPS buffer without EDTA (50 mM MOPS, 50 mM Tris base, 0.1% [w/v] SDS, pH 7.7) supplemented with 2.5 mM sodium bisulfite, for 2 h at 150 V constant. Protein phosphorylation pattern was visualized by Coomassie staining.

**Western blotting and antibodies**

For western blotting, samples were resolved by SDS-PAGE on NuPAGE 4-12% bis-tris gel using MES buffer followed by transfer onto 0.2 µm PVDF membranes using a Bio-Rad Trans-Blot Turbo semi-dry system. For blotting of Phos-Tag gels, samples were resolved by SDS-PAGE as described above and the gel was incubated in Tris-glycine running buffer with EDTA (25 mM Tris base, 192 mM glycine, 0.1% [w/v] SDS, pH 8.3 supplemented with 50 mM EDTA, pH 7.4) for 20 min at RT prior to the transfer step. Membranes were blocked in 5% (w/v) BSA in TBS with 0.1% Tween-20 (TBS-T; 20 mM Tris-HCl, pH 7.4, 150 mM NaCl, 0.1% [v/v] Tween-20) for 30 min at RT, rinsed with TBS-T and incubated in primary antibodies overnight at 4°C. Membranes were then washed 3 x 10 min in TBS-T and incubated with HRP-conjugated secondary antibodies for 1 h at RT. After a further 3 x 10 min washes in TBS-T, blots were developed by chemiluminescence using SuperSignal Chemiluminescent Substrate (Thermo Fisher) on a Bio-Rad ChemiDoc MP imaging system.

Antibodies dilutions were made in 3% (w/v) BSA in TBS-T. Rabbit anti-AKT (1:2000, #MA5-51195) and rabbit anti-SIN1-pS510 (1:1000, PA5-105694) were purchased from Invitrogen (Thermo Fisher). Rabbit anti-AKT-pT308 (1:1000, #2965) and rabbit anti-AKT-pS473 (1:2000, #4060) were purchased from Cell Signaling Technologies. Rabbit anti-SIN1 (1:1000, 07-2276-I) and rabbit anti-SIN1-pT86 (1:1000, ABS1519) were purchased from Merck.

**Cryo-ET sample preparation**

For mTORC2 immobilization on LUVs, 4 µM mTORC2 was mixed with 1 mg/mL 100 nM 5% PIP2 LUVs in a total volume of 150 µL made up in buffer (25 mM HEPES, pH 7.5, 150 mM NaCl, 0.5 mM $MgCl_2$, 0.5 mM TCEP) and incubated for 15 min at RT. LUVs were generated as described in the above method and used fresh. The sample was centrifuged for 1 min at 4,000 x *g* to remove large aggregates of LUVs and the supernatant carefully moved to a fresh 1.5 mL tube. LUVs were then pelleted by centrifugation at 10,000 x *g* for 10 min at 12°C. The supernatant was discarded and the mTORC2-LUV pellet rinsed twice with 100 µL buffer, being careful not to disturb the pellet. The pellet was then resuspended in 10 µL of buffer, followed by incubation in a water bath sonicator, for 15 min at 4°C to ensure thorough resuspension of the sample. The sample was diluted 3:1 (sample:buffer) and then 10 nm fiducial gold added at a ratio of 8:1 (sample:fiducial). Quantifoil R2/2 Au200 grids were glow-discharged for 1 min using a S150B Sputter Coater (Edwards) at 40 mA. Sample grids were prepared using 3.3 µL of sample, blotted for 4 s with a blot force of 10, then immediately plunge-frozen in liquid ethane (-180°C) on a Vitrobot Mark IV plunger (Thermo Fisher) equilibrated to 18°C, 100% humidity.

**Cryo-ET data collection and analysis**

The Cryo-ET dataset was collected on a 300 kV Titan Krios G4 with a Falcon 4i direct electron detector and a Selectris-X energy filter (all Thermo Fisher) using a 10 eV slit width. Movies were recorded at a magnification of 81,000x, corresponding to a pixel size of 1.514 Å/pixel. Tilt-series were collected in EER format at 3° increments in a dose-symmetric scheme from -60° to +60° using PACE-tomo scripts within SerialEM (*56*) and a dose of 3.4 $e^{-}$/Å$^{2}$ per tilt (140 $e^{-}$/Å$^{2}$ total dose). A total of 180 tilt-series were collected.

Tomography processing was performed using the RELION 5.0 tomography pipeline (*57*) and is outlined in **fig. S3**. Tilt-series were imported, movies aligned using MOTIONCOR2 with EER fractionation of 0.5 $e^{-}$/Å$^{2}$ per movie fraction and contrast transfer function (CTF) estimation performed using CTFFIND4 (*58*). Tilt-series were aligned by fiducial-based alignment using IMOD within RELION 5 and tomograms reconstructed at 10 Å/pixel for particle picking. Particles were picked using SPHIRE-crYOLO tomography picking mode (*59*), using a model trained on manually picked coordinates from 12 tomograms (**fig. S3A**). Processing was performed using an un-binned particle box size of 512 x 512 pixels and a cropped box size of 352 x 252 pixels (~530 Å) during sub-tomogram particle extraction, and a soft-spherical mask of 320 Å was used during refinements. A *de novo* 3D initial reference was generated and used as a low-pass filtered (40 Å) reference for un-supervised 3D classification. The dataset was split into 2 equal sub-sets which were processed in parallel classification jobs. 3D classification was performed in bin6 (9.084 Å/pixel) for a total of 4 iterations. After each iteration, 3D classes were inspected in ChimeraX and classes containing membrane-bound mTORC2 were grouped for further processing. 3D classification also revealed classes which contained mTORC2 in the absence of membrane. Further classification of these non-membrane particles revealed further mTORC2 membrane-bound particles (**fig. S3B**) which we grouped back into the final sub-tomogram set. The final set of 38,519 sub-tomogram particles underwent 3D auto-refinement first in bin6, followed by iterative extraction and refinement at lower binning factors (bin4, 2, and 1). To improve mTORC2 particle alignment, we performed focused refinement using a mask around the mTORC2 protein only, excluding the noisy membrane signal. After focused refinement and 3 rounds of iterative CTF refinement and Bayesian polishing, the resolution of membrane-bound mTORC2 was 6.4 Å. This focused mTORC2 map was used for model building. An mTORC2 dimer model was generated using the SPA autophosphorylated mTORC2 dimer as a starting model, with the mTORC2 N-HEAT built using the human mTOR AlphaFold model from the Alpha Fold Protein Structure Database (ID: AF-P42345-F1-v4). Model geometry and fit was further improved using Coot (*60*) and ISOLDE (*61*) in ChimeraX, before final real-space refinement in Phenix (*62*). Final refinement and model statistics (**Table S2**) were calculated using globally sharpened mTORC2 protein focused experimental maps from RELION postprocessing and these maps were submitted as the primary map during PDB/EMDB deposition.

**Cryo-EM sample preparation**

For dephosphorylated mTORC2, a 50 µL frozen aliquot of dephospho-mTORC2 (prepared as described above) was thawed, cleared by centrifugation at 20,000 x *g* for 2 min at 4°C and loaded on a Superose 6 Increase 3.2/300 analytical column (Cytiva) equilibrated in SEC buffer (20 mM HEPES, pH 7.5, 250 mM NaCl, 1 mM TCEP). The peak fraction was diluted 1.9 µM (1.1 mg/mL) mTORC2 for grid preparation. For autophosphorylated-mTORC2, dephospho-mTORC2 was thawed and incubated for 20 min at 30°C in a 60 µL reaction made up in SEC buffer to a final concentration of 10 µM mTORC2, 1 mM ATP, 10 mM $MgCl_2$. The autophosphorylated mTORC2 was then purified by SEC and grids prepared as described for dephospho-mTORC2. The peak fraction was diluted to 1.5 µM (0.9 mg/mL) mTORC2 for grid preparation.

For AKT-phosphorylated mTORC2, untreated mTORC2 was first dephosphorylated with λ-PP for 1 h at 30°C in a 60 µL reaction made up in SEC buffer to a final concentration of 9 µM mTORC2, 1.6 µM λ-PP, 1 mM $MnCl_2$. The dephosphorylated mTORC2 was isolated by SEC as described above, then phosphorylated with $AKT^{WT}$ for 1 h at 30°C in a 25 µL reaction made up in SEC buffer to a final concentration of 1.5 µM mTORC2, 1 µM $AKT^{WT}$, 1 mM ATP, 10 mM $MgCl_2$. This reaction was used directly for grid preparation.

Cryo-EM grids for all final data collection samples were prepared using the same protocol. UltrAuFoil R1.2/1.3 Au300 grids were glow-discharged for 80 s using a S150B Sputter Coater (Edwards) at 40 mA. Sample grids were prepared using 3.3 µL of sample, blotted for 3.5 s with a blot force of 15, then immediately plunge-frozen in liquid ethane (-180°C) on a Vitrobot Mark IV plunger (Thermo Fisher) equilibrated to 14°C, 100% humidity.

**Cryo-EM data collection and processing**

All 3 SPA datasets were collected on a 300 kV Titan Krios with a Falcon 4i direct electron detector and a Selectris-X energy filter (all Thermo Fisher) using a 10 eV slit width. Images were recorded using automated data collection in EPU (Thermo Fisher) at a magnification of 130,000x, corresponding to a pixel size of 0.955 Å/pixel. Images were collected in electron counting mode using a defocus range of -1 to -3.0 µm. A total dose of ~45, 50 and 50 $e^-/Å^2$ was fractionated into 42, 45 and 50 movie frames for dephos, auto- and AKT-phosphorylated mTORC2, respectively. A total of 9,381, 9,786 and 7,153 micrographs were collected for dephospho-, auto- and AKT-phosphorylated mTORC2, respectively.

Processing of all datasets was performed using RELION 5.0 (*63*) and is outlined in **fig. S6**. The images were imported, aligned using MOTIONCOR2 and contrast transfer function (CTF) estimation performed using CTFFIND4 (*58*). Particles were picked using SPHIRE-crYOLO (*59*), utilizing a model trained on manually-picked particle coordinates from 10 micrographs in the phospho-mTORC2 dataset; this model was used for all 3 datasets. A second sets of particles were picked using RELION's reference-based template-matching autopicker, utilizing three 2D-references generated by 2D classification of ~4,000 manually picked particles as templates, this was performed independently for each dataset. For processing, bin4 down-sampling was utilized for 2D & 3D classification; the un-binned particle box size was 512 x 512 pixels and a 360 Å soft spherical mask was utilized throughout. A single round of reference-free 2D classification was performed for each picking method, before combining mTORC2 classes and removing duplicate particles within RELION. The combined particles were then subjected to 2 further rounds of 2D classification. A *de novo* 3D initial reference was generated, which was then used as a 40 Å low-pass filtered reference for un-supervised 3D classification into 4 classes. This generally yielded a single mTORC2 class with high-resolution features which was carried forward. Particles were re-extracted and underwent 3D auto-refinement in bin2 followed by bin1, using no mask around the mTORC2 dimer. Refined dimer particles then underwent a single round of CTF refinement followed by Bayesian polishing in RELION, with further 3D auto-refinement (using Blush regularization) and post-process B-factor sharpening yielding reconstructions of 3.3, 3.1 and 3.0 Å global resolution for dephospho-, auto- and AKT-phosphorylated mTORC2, respectively. Commonly across all 3 datasets, local resolution estimates for the consensus dimer maps showed higher resolution in a single half of the dimer. Focused 3D refinement using a mask around the better half of the dimer resulted in increased global resolutions of 3.0, 2.8 and 2.6 Å for dephospho-, auto- and AKT-phosphorylated mTORC2 datasets, respectively. Despite apparent asymmetry in map quality of each protomer within the dimer consensus refinements, further focused refinements of the second, lower quality, protomer produced well aligned maps (3.0, 3.1 and 2.6 Å resolution for dephospho-, auto- and AKT-phosphorylated mTORC2, respectively) suggesting the asymmetry is derived from flexibility between the dimer halves, rather than any heterogeneity within each individual protomer. Focused refinements for each protomer were aligned to the dimer consensus map and combined in ChimeraX to generate a symmetric, high-resolution dimer composite map for each mTORC2 state.

Composite dimer and focused protomer maps were sharpened using cryoTEN (*64*) and used for model building. The cryoTEN sharpened maps were used to show EM density in main text figures. An initial high-resolution protomer model was built into the autophosphorylated mTORC2 focused protomer map using ModelAngelo in RELION (*65*). Lower resolution regions which were not built fully by ModelAngelo were modelled using predicted models from the AlphaFold database, with manual alteration and building in Coot (*60*). Lower resolution regions which were not built fully by ModelAngelo were modelled using predicted models from the AlphaFold database, with manual fitting in Coot (*60*). Model geometry and fit was further improved using ISOLDE (*61*) in ChimeraX, before final real-space refinement in Phenix (*62*). The auto-mTORC2 refined protomer model was used as the starting model for all other models which were similarly built in Coot and ISOLDE before Phenix real-space refinement. For dimer models, two copies of the corresponding protomer model were fit into composite dimer map followed by building and refinement as described for the protomer models. For all models, final refinement and model statistics (**Table S3**) were calculated using globally sharpened consensus maps from RELION postprocessing. Gold-standard FSC and final resolution for all datasets (**fig. S7**) were calculated using the 3DFSC server (3dfsc.salk.edu; (*66*)).

**Differential scanning fluorimetry (DSF)**

For DSF measurements, purified proteins were defrosted and diluted to 1 mg/mL in kinase buffer (25 mM HEPES, pH 7.5, 100 mM NaCl, 5% (v/v) glycerol, 0.5 mM TCEP). Thermal stability was assessed by monitoring internal tryptophan fluorescence of 10 µL samples on a Prometheus NT.48 (NanoTemper). Fluorescence at 330 and 350 nm wavelength was recorded across a 15-95°C gradient at 2°C/min and melting temperatures were determined using PR.ThermControl software (NanoTemper). Melting temperatures were derived from the inflection point of plotting the ratio of fluorescence at 350/330 nm versus temperature (i.e. the temperature at which 50% of the protein is unfolded). Final melting temperatures are reported as the mean ± SD of 3 capillary samples per protein.

**Hydrogen-deuterium exchange mass spectrometry (HDX-MS)**

A preincubation was performed by adding dilution buffer (50 mM Bicine pH8.5, 300mM NaCl, 0.5 mM TCEP) to yield solutions that were 5.5 µM of dephosphorylated, AKT-phosphorylated or autophosphorylated mTORC2 complex. Samples were incubated for 15 minutes on ice and then exposed for different deuteration times (3 s on ice, 3, 30, 300 and 3000 s at room temperature) to a deuterated buffer: 50 mM Bicine pD8.5, 300 mM NaCl, 0.5mM TCEP with $D_2O$ (Thermo Scientific, 166300100) to yield a final concentration 0.3 µM of mTORC2 samples in 95% $D_2O$. Experiments were performed in triplicate for each timepoint and exchange reactions were quenched in 2 M Guanidine-Hydrochloride, 100 mM glycine, formic acid (Fisher Chemical, 10596814), LC-MS grade water (Romil, H949), pH 2.0 (final pH was 2.5). Samples were flash-frozen in $LN_2$ immediately and then stored at -80°C until analysis. Prior to LC-MS analysis, samples were quickly thawed and injected on a HDX Manager coupled to an Acquity UPLC M-Class system (Waters) set at 0.1 °C. Samples were then digested at 15 °C using a 20 mm x 2.0 mm home-made pepsin column (Pepsin from Thermo Scientific, 20343 and hardware from Upchurch Scientific, C130-B) and loaded on a UPLC pre-column (ACQUITY UPLC BEH C18 VanGuard pre-column, 2.1 mm I.D. x 5 mm, 1.7 µM particle diameter, Waters, 186003975) for 2 min at 100 µL/min. Digested peptides were then eluted over 20 min from the pre-column onto an ACQUITY UPLC BEH C18 column (1.0 mm I.D. x 100 mm, 1.7 µM particle diameter, Waters, 186002346) using a 5-43% gradient of Acetonitrile (Romil, H050), 0.1% formic acid and interfaced to a Q Exactive™ HF-X hybrid quadrupole-Orbitrap mass spectrometer (Thermo Scientific, USA) *via* a Heated-electrospray ionization source (HESI). For the undeuterated analysis, MS/MS data were acquired in data dependent mode with a Top-20 method as follows: full mass scans were acquired (R = 60,000; 400-800 m/z range; max IT = 200 ms, AGC Target = $3e^6$) followed by MS/MS scans using higher energy collision dissociation (HCD) with normalized collision energy at 30% (R = 60,000;

isolation window: 1.2m/z; max IT = 100 ms; AGC Target = $5e^5$; Charge exclusion: 6-8 and >8; Dynamic exclusion = 30 s). For deuterated analysis, MS1 only data were acquired as follows: R = 120,000; 400-800 m/z range; max IT = 200 ms and AGC Target = $3e^6$. The instrument was calibrated in positive mode using a Pierce™ LTQ ESI Positive Ion Calibration Solution (Thermo Scientific, 88322). Peptide identification was performed with undeuterated samples using Proteome Discoverer v3.2.0.450 (Thermo Scientific) using SEQUEST HT with a home-made protein sequence library containing mTOR, Rictor, SIN1, mLST8 and pepsin sequences. Peptide length was set to a minimum of 5 residues and a maximum of 25 residues. Oxidation of methionine, acetylation of protein N-term, phosphorylation of serine, threonine and tyrosine residues, loss of methionine and loss methionine + acetylation were set as variable modifications. Data were extracted using peptide mass tolerance of 5 ppm and MS/MS tolerance at 0.02 Da with a false discovery rate of 1% at precursor and fragment levels. Deuterium uptakes for all identified non-phosphorylated peptides were then calculated using HDExaminer 3.4.0 (Trajan Scientific and Medical) with an initial automated spectral processing step followed by a manual inspection of individual peptides for sufficient quality where only one charge state per peptide was kept. Deuterium uptakes were not corrected for back-exchange and are reported as relative. HDX-MS results were statistically validated using an in-house program (Archaeopteryx), where the *p*-value threshold was set to 0.05 for the *t*-test. Only statistically significant peptides showing a difference greater than 0.3 Da and greater than 5% were kept. HDX-MS results were exported on AKT-phosphorylated and auto-phosphorylated models using PyMOL 2.5.4 (www.pymol.org). The mass spectrometry data have been deposited to the ProteomeXchange Consortium via the PRIDE partner repository (*67*) with the dataset identifier PXD071935.

### Supplementary Figures

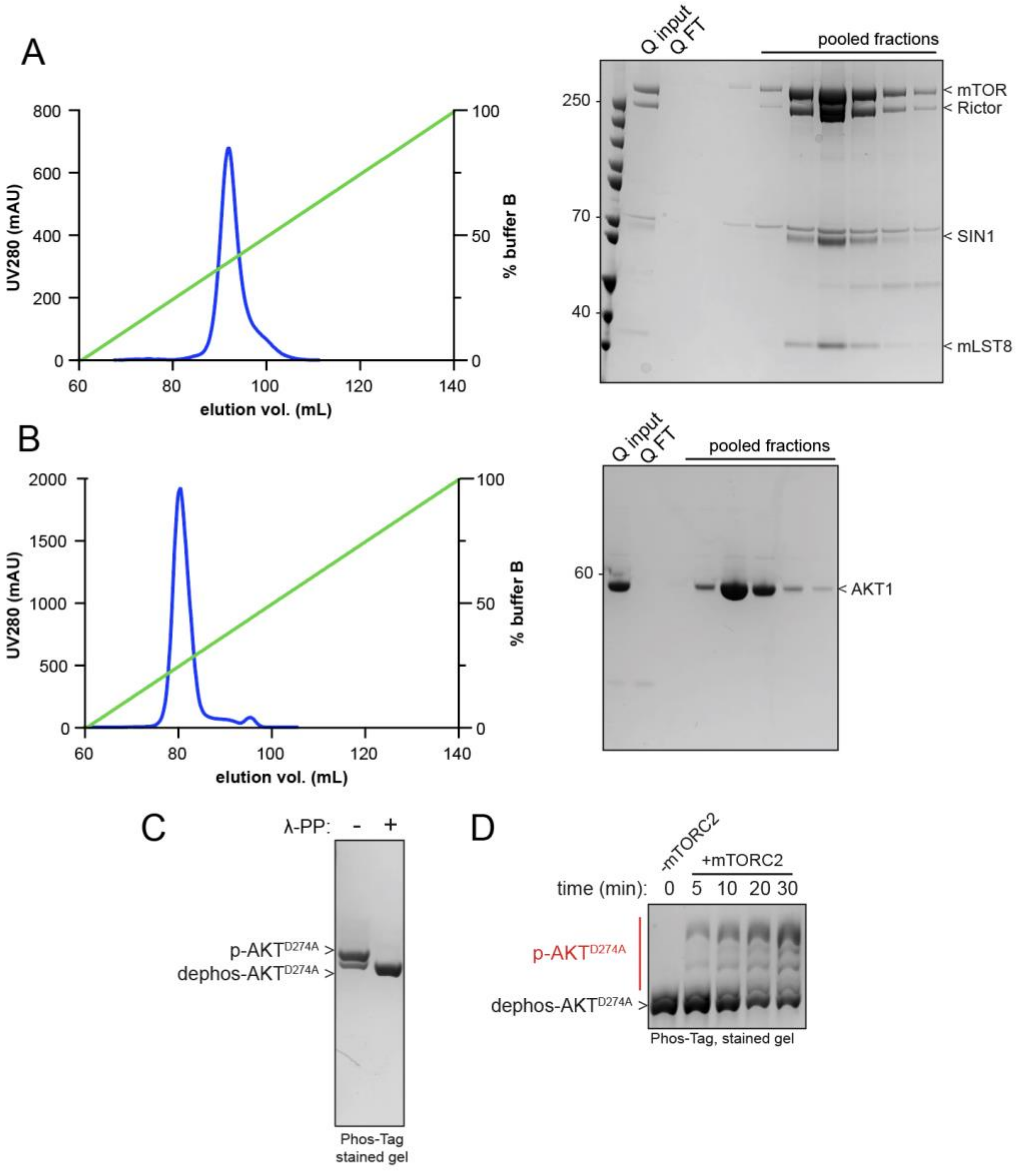

**Fig. S1. Purification and characterization of mTORC2 and AKT$^{D274A}$ substrate for kinase assays.**

(A) Representative preparation of mTORC2. Left: UV absorbance at 280 nm (UV280) trace for anion exchange chromatography (AEX) of 2xStrep-tagged mTORC2 on a HiTrap Q HP 5 mL column (Q) after Strep-tag affinity purification. Right: SDS-PAGE Coomassie-stained gel of peak fractions from the AEX step. Fractions pooled for further analysis are indicated.

(B) Representative purification of AKT$^{D274A}$. Left: UV280 trace for AEX of AKT$^{D274A}$ after Strep-tag affinity purification followed by overnight TEV cleavage and dephosphorylation with lambda protein phosphatase ($\lambda$-PP). Right: SDS-PAGE Coomassie-stained gel of peak fractions from the AEX step. Fractions pooled for further analysis are indicated.

(C) Phos-Tag SDS-PAGE analysis of purified $AKT^{D274A}$ before (-) and after (+) overnight treatment with $\lambda$-PP. $AKT^{D274A}$ as purified from Expi293F cells has a single phosphorylated band presumed to be co-translational phosphorylation at T450.

(D) mTORC2 activity assay using purified mTORC2 (0.2 µM) with 25 µM dephosphorylated $AKT^{D274A}$ as a model substrate. Reaction was carried out for the indicated timepoints at 30°C with 1 mM ATP, 10 mM $MgCl_2$.

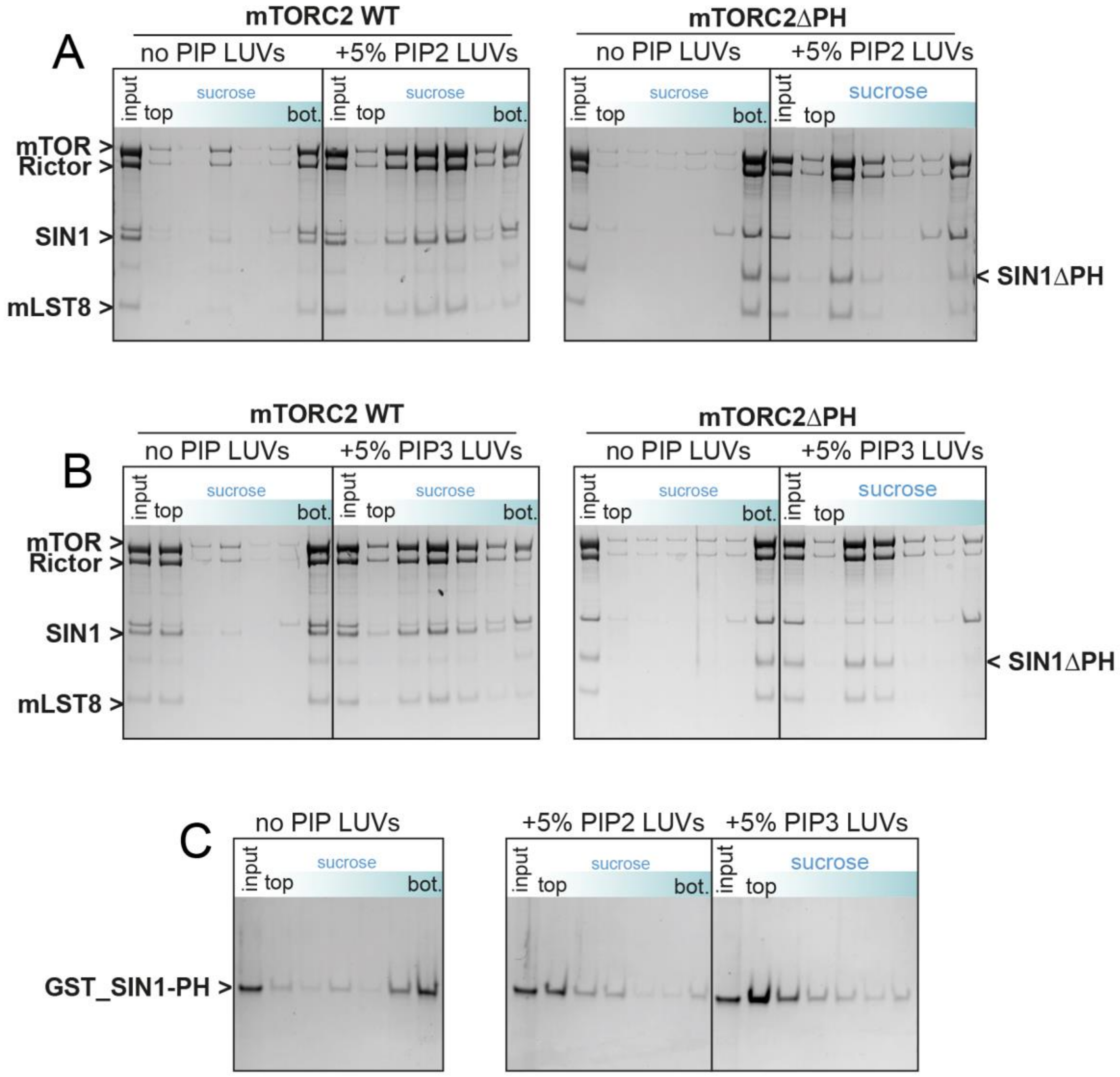

**Fig. S2. Characterization of SIN1-PH in membrane binding.**

(A) Liposome flotation assays of mTORC2 WT or SIN1ΔPH (mTORC2ΔPH). mTORC2 in the presence of no PIP (base lipids only) or 5% PIP2 containing 100 nm LUVs were analyzed in a 35% (bottom) to 25% (top) sucrose gradient. The mTORC2/lipid mixture was placed at the bottom of the gradient, with liposomes and bound protein floating into top fractions during centrifugation.

(B) Liposome flotation assays as described in (A), using PIP3 100 nm LUVs.

(C) Liposome flotation assays of GST-tagged SIN1-PH domain in the presence of 0% PIP, 5% PIP2 or 5% PIP3 containing 100 nm LUVs, performed as described in (A).

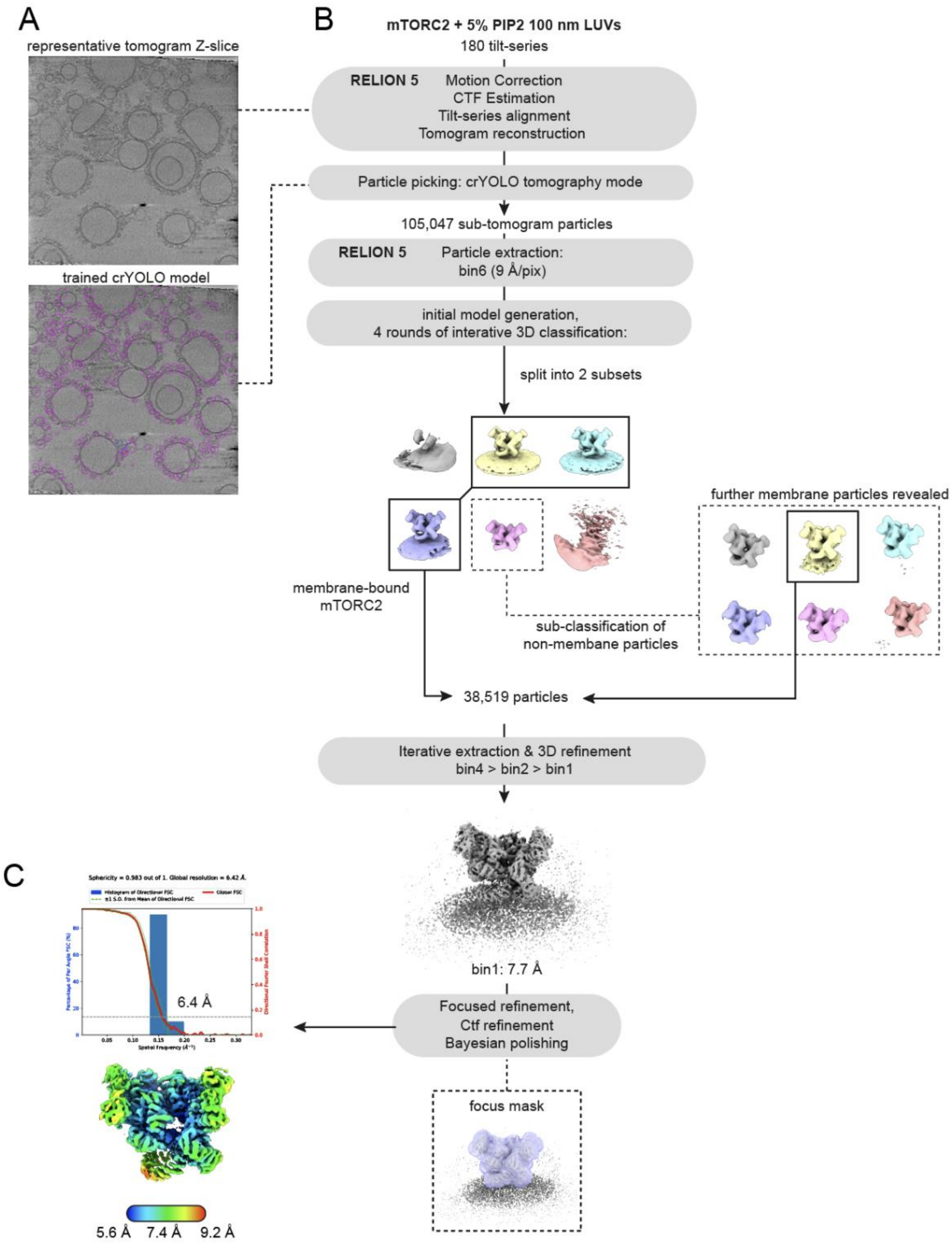

**Fig. S3. Cryo-ET sub-tomogram averaging strategy for membrane-bound mTORC2.**

(A) Top: Representative Z-slice of a reconstructed tomogram, binned to 10 Å/pixel. Bottom: Example of picked sub-tomogram particles (magenta circles) using a crYOLO model trained on 12 manually picked tomograms.

(B) Tomogram reconstruction and sub-tomogram averaging workflow.

(C) Directional FSC plot for the mTORC2 membrane-bound map, generated using the 3DFSC server (3dfsc.salk.edu; (*66*)). The map is shown below, colored by local resolution estimates calculated in RELION 5.

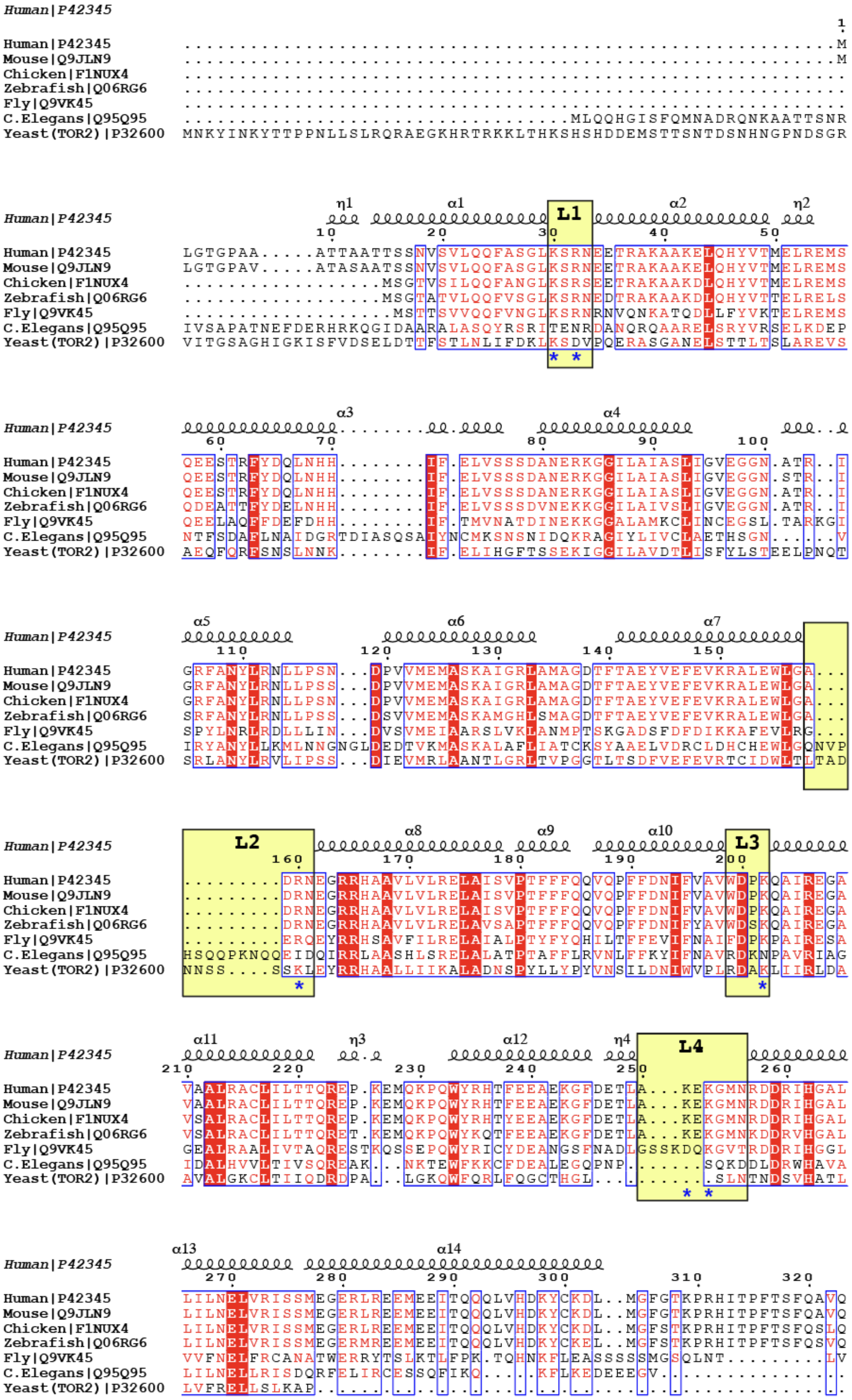
Human|P42345
Mouse|Q9JLN9
Chicken|F1NUX4
Zebrafish|Q06RG6
Fly|Q9VK45
C.Elegans|Q95Q95
Yeast(TOR2)|P32600
η1 α1 L1 α2 η2
α3 α4
α5 α6 α7
L2 α8 α9 α10 L3
α11 η3 α12 η4 L4
α13 α14

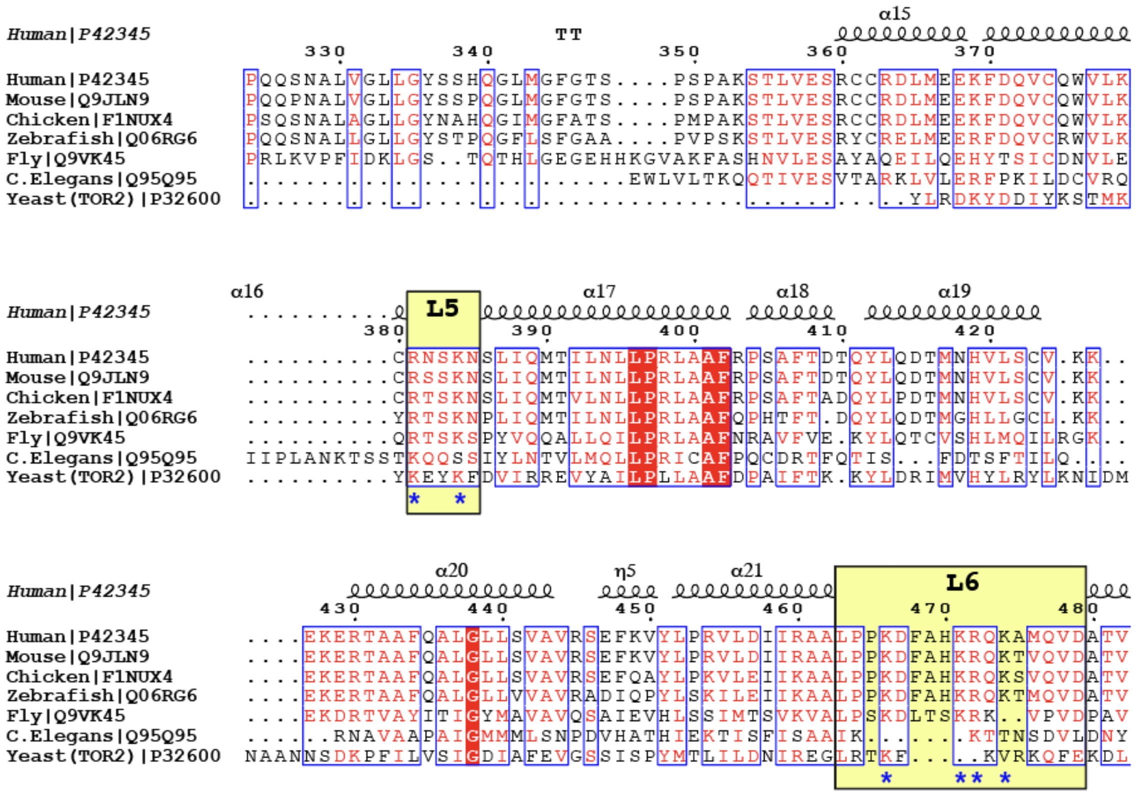

**Fig. S4. Multiple sequence alignment of residues 1-482 of the mTOR N-HEAT domain.** Sequences are numbered as for human mTOR. Membrane proximal loops (L1-L6) from main text Fig. 1E are highlighted by yellow boxes. Basic residues within these loops are marked with a blue asterisk. The annotated secondary structure was derived from the AlphaFold model of human mTOR from the Alpha Fold Protein Structure Database (ID: AF-P42345-F1-v4, retrieved October 2025). Invariant residues are shown as white letters on a red background. Blue frames highlight columns in which 70% of residues share similar physicochemical properties (red residues) across species. Alignment was performed using Clustal Omega (*68*) and graphics generated in ESPript 3.0 (https://endscript.ibcp.fr, (*69*)). Sequences were retrieved from Uniprot and Uniprot IDs are shown next to species name.

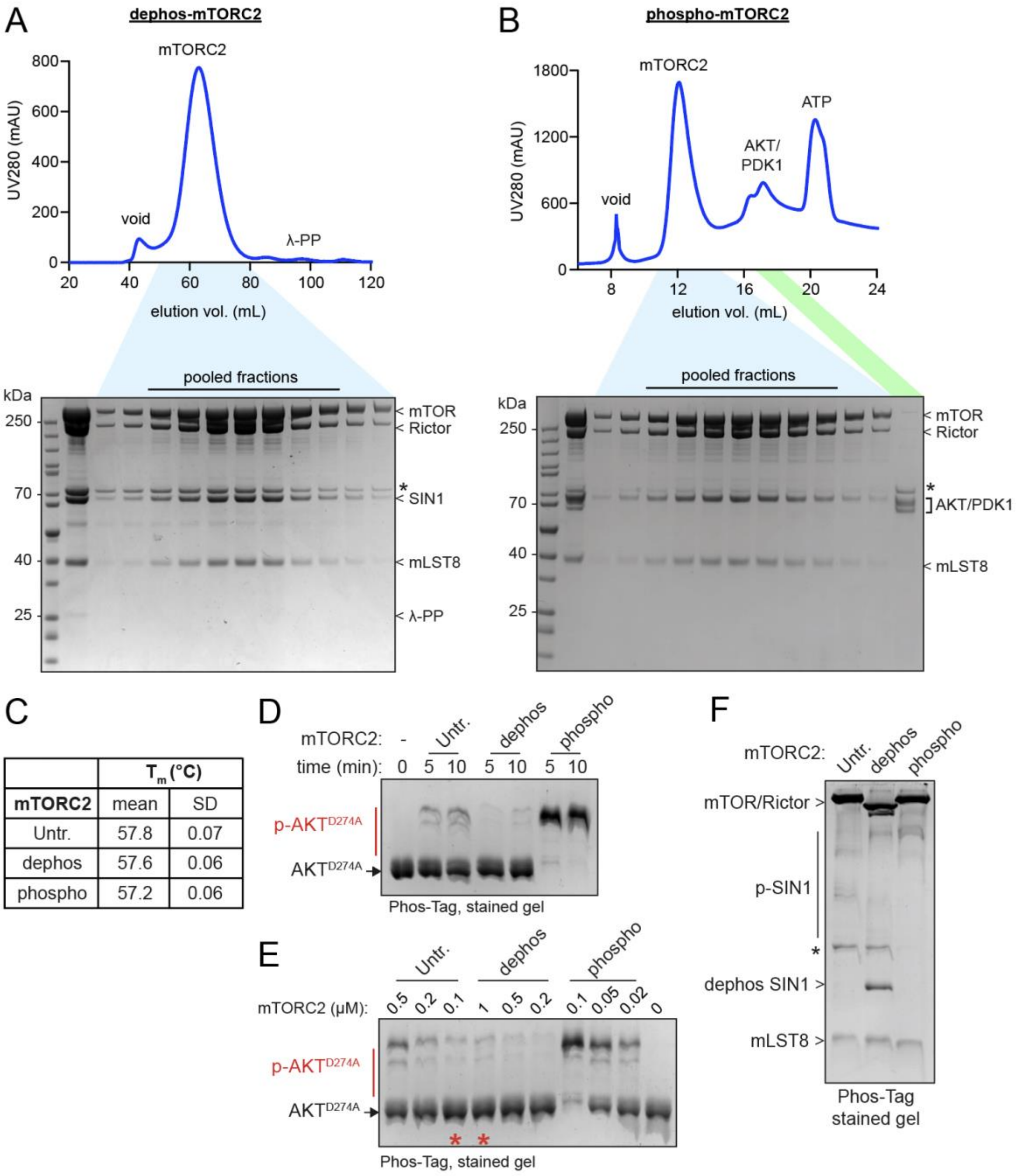

**Fig. S5. Purification of dephosphorylated and AKT-phosphorylated mTORC2.**

(A) Representative purification of dephosphorylated mTORC2 (dephos-mTORC2). Top: UV absorbance at 280 nm (UV280) trace for size-exclusion chromatography (SEC) of dephos-mTORC2 on a Superose 6 16/600 column after overnight dephosphorylation with lambda protein phosphatase (λ-PP). Below: SDS-PAGE Coomassie-stained gel of peak fractions from the SEC step. The 25 kDa λ-PP is visible in the SEC input and is removed during the chromatography step.

(B) Representative purification of AKT-phosphorylated mTORC2 (phospho-mTORC2). Top: UV280 trace for SEC of phospho-mTORC2 on a Superose 6 Increase 10/300 column after overnight dephosphorylation with λ-PP, followed by *in vitro* phosphorylation with AKT (5 µM AKT was pre-phosphorylated with 1 µM PDK1, then used at an equimolar mTORC2:AKT ratio for *in vitro* phosphorylation). Below: SDS-PAGE Coomassie-stained gel of peak fractions from the SEC step. AKT, PDK1 and ATP are isolated from the phospho-mTORC2 during the chromatography step. (*) denotes HSP70 contaminant, which is removed by the ATP during the phospho-mTORC2 preparation.

(C) Global thermostability of purified untreated (Untr.), dephospho- and AKT-phospho-mTORC2 proteins. Melting temperature (Tm) for each complex was determined by nanoDSF at 1 mg/mL mTORC2. Data represent mean ± SD of n = 3 technical replicates.

(D) Confirming activation/inhibition of mTORC2 protein preps. mTORC2 activity assays of Untr., dephospho- and AKT-phospho-mTORC2 (0.1 µM) incubated for the indicated time with 25 µM $AKT^{D274A}$ substrate at 30°C, 1 mM ATP, 10 mM $MgCl_2$. Dephosphorylation inactivates mTORC2 whilst AKT phosphorylation activates when compared to Untr-mTORC2.

(E) mTORC2 activity assays of untreated, dephospho- and AKT-phospho-mTORC2 at varying enzyme concentrations. The red asterisks highlight that 1 µM dephospho-mTORC2 produces similar levels of $AKT^{D274A}$ substrate phosphorylation as 0.1 µM of untreated mTORC2.

(F) Phos-Tag SDS-PAGE analysis of Untr., dephospho- and AKT-phospho-mTORC2 protein preps. SIN1 is completely dephosphorylated in dephos- and subsequently fully re-phosphorylated in phospho-mTORC2.

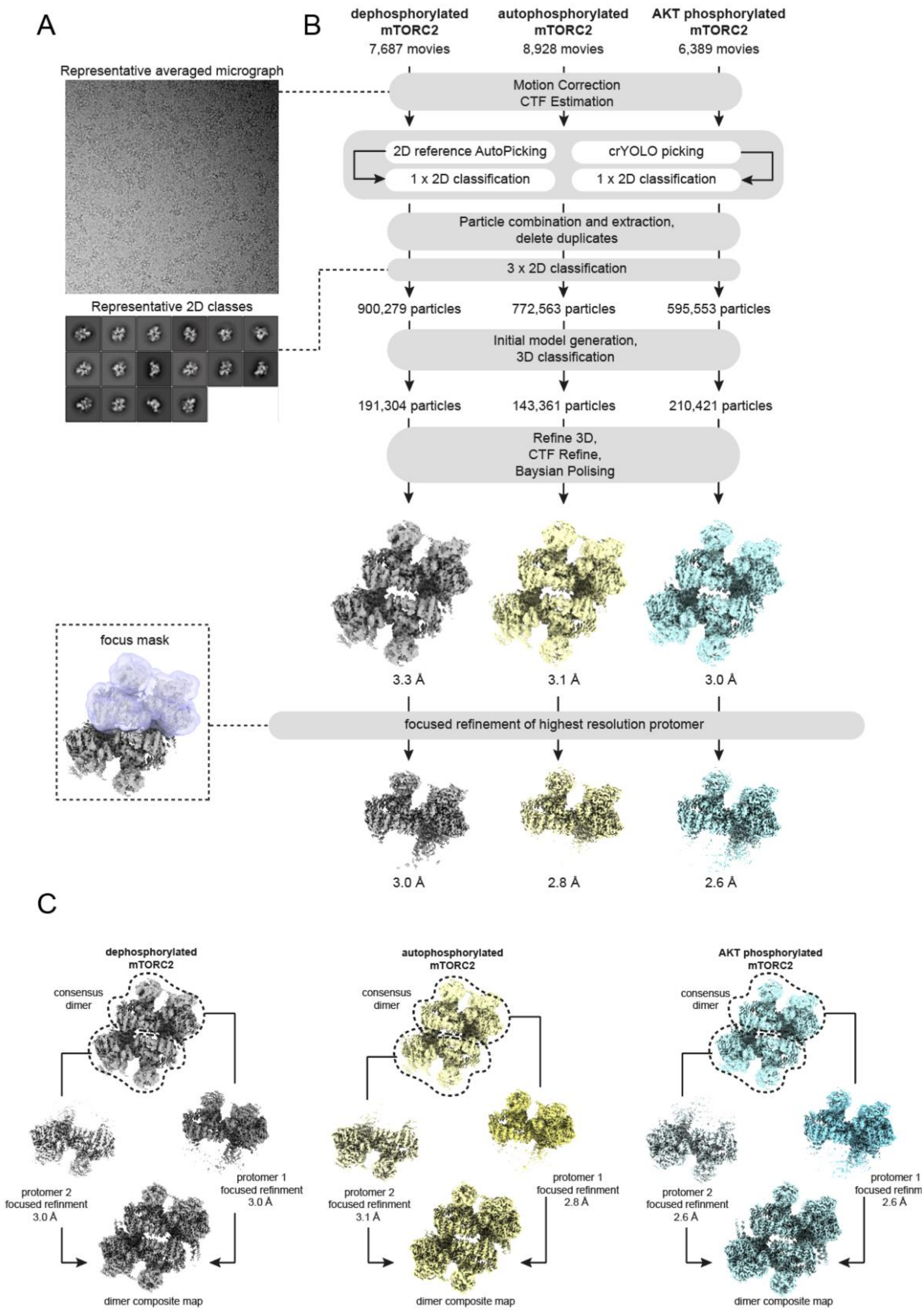

**Fig. S6. Cryo-EM single particle analysis (SPA) of mTORC2 phosphorylation.**

(A) Representative micrograph and 2D classes of mTORC2, taken from the AKT phosphorylated dataset.

(B) SPA data processing workflow – datasets were analyzed independently, but using a common strategy.

(C) Dimer composite map generation. Despite apparent asymmetry in resolution between protomers in the consensus dimer maps (see local resolution in fig. S7), focused refinement of individual protomers yielded high quality maps. Protomers were focus refined individually, aligned to the consensus dimer map and used to generate a composite dimer map generated in ChimeraX. These maps were used for mTORC2 dimer model building.

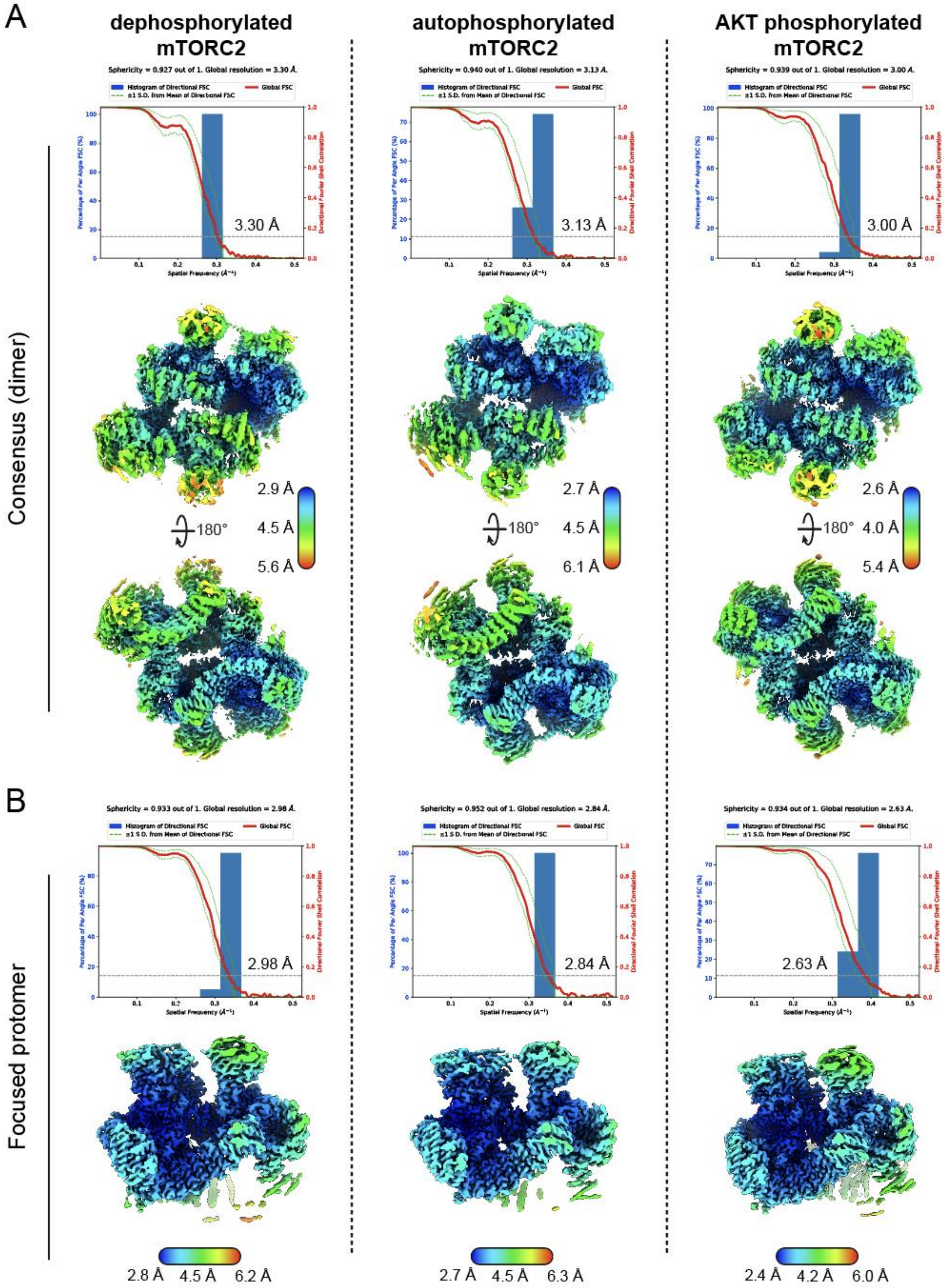

**Fig. S7. Cryo-EM map quality metrics for mTORC2 phosphorylated states.**

(A) Directional FSC plots for the consensus dimer maps of dephospho-, auto- and AKT-phospho-mTORC2, generated using the 3DFSC server (3dfsc.salk.edu; (*66*)). Top and bottom views of the mTORC2 dimer maps are shown, colored by local resolution estimates calculated in RELION 5.

(B) Metrics for the focused protomer maps, as described for dimers in (A).

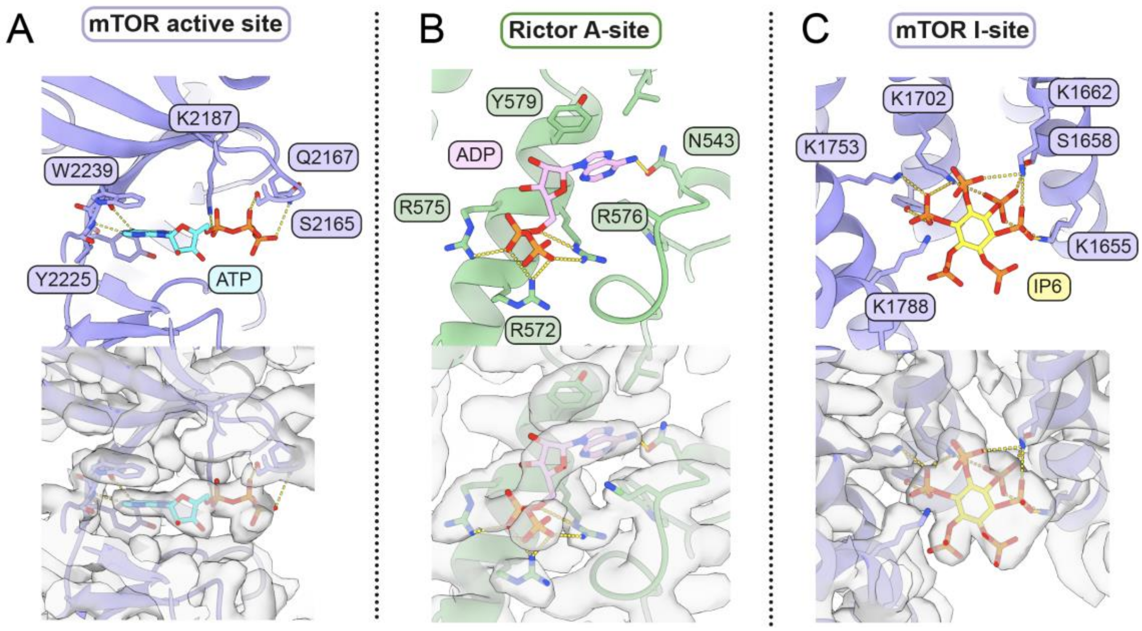

**Fig. S8. Ligand binding in AKT-phosphorylated mTORC2.**

(A-C) For all panels, the atomic model is shown above, with key residues involved in ligand binding labelled and shown as sticks. H-bonds are shown as yellow dashes. The sharpened cryo-EM density map for each ligand is shown below.

(A) ATP (cyan sticks) is shown bound in the active site of mTOR (slate blue).

(B) ADP (pink sticks) is shown bound in the A-site of the Rictor HEAT domain (green).

(C) Inositol hexaphosphate (IP6; yellow sticks) is shown bound in the I-site within the mTOR FAT domain (slate blue). This panel shows IP6 bound in the phospho-mTORC2 model, but IP6 was present in all three mTORC2 states.

A

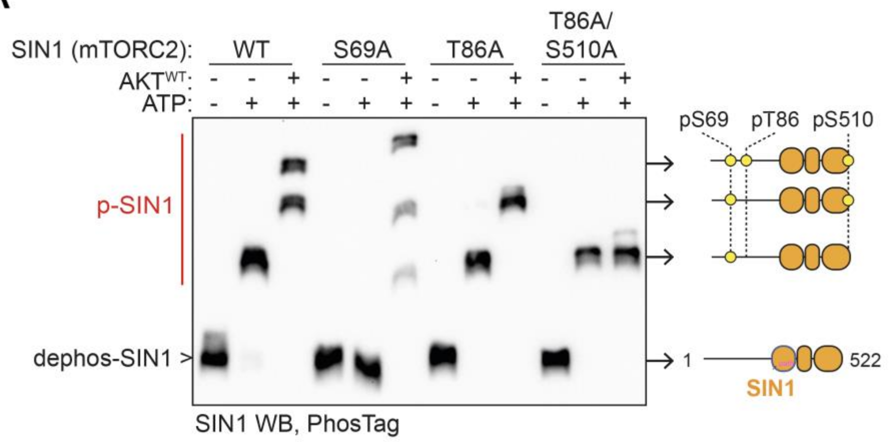

B

| | $T_m$ (°C) | |
|---|---|---|
| **mTORC2:** | mean | SD |
| SIN1-S69A | 57.7 | 0.07 |
| SIN1-T86A | 57.5 | 0.10 |
| SIN1-T86A/S510A | 57.2 | 0.06 |

**Fig. S9. Analysis of SIN1 phosphorylation and phospho-site mutants.**

(A) PhosTag SDS-PAGE western blotting of SIN1 after treatment of 1 µM dephosphorylated mTORC2 SIN1 WT and mutants with 1 mM ATP ± 1 µM AKT$^{WT}$ for 20 min at 30°C. S69A mutation removes autophosphorylation upon ATP treatment, showing this to be the SIN1 autophosphorylation site. Tandem mutation of T86A/S510A produces only autophosphorylation, even in the presence of AKT$^{WT}$, showing that T86 and S510A are the only AKT phosphorylation sites within SIN1.

(B) Global thermostability of purified dephosphorylated mTORC2 complexes containing SIN1 phospho-site mutants. The melting temperature (Tm) for each complex was determined by nanoDSF at 1 mg/mL mTORC2. Data represent mean ± SD of n = 3 technical replicates.

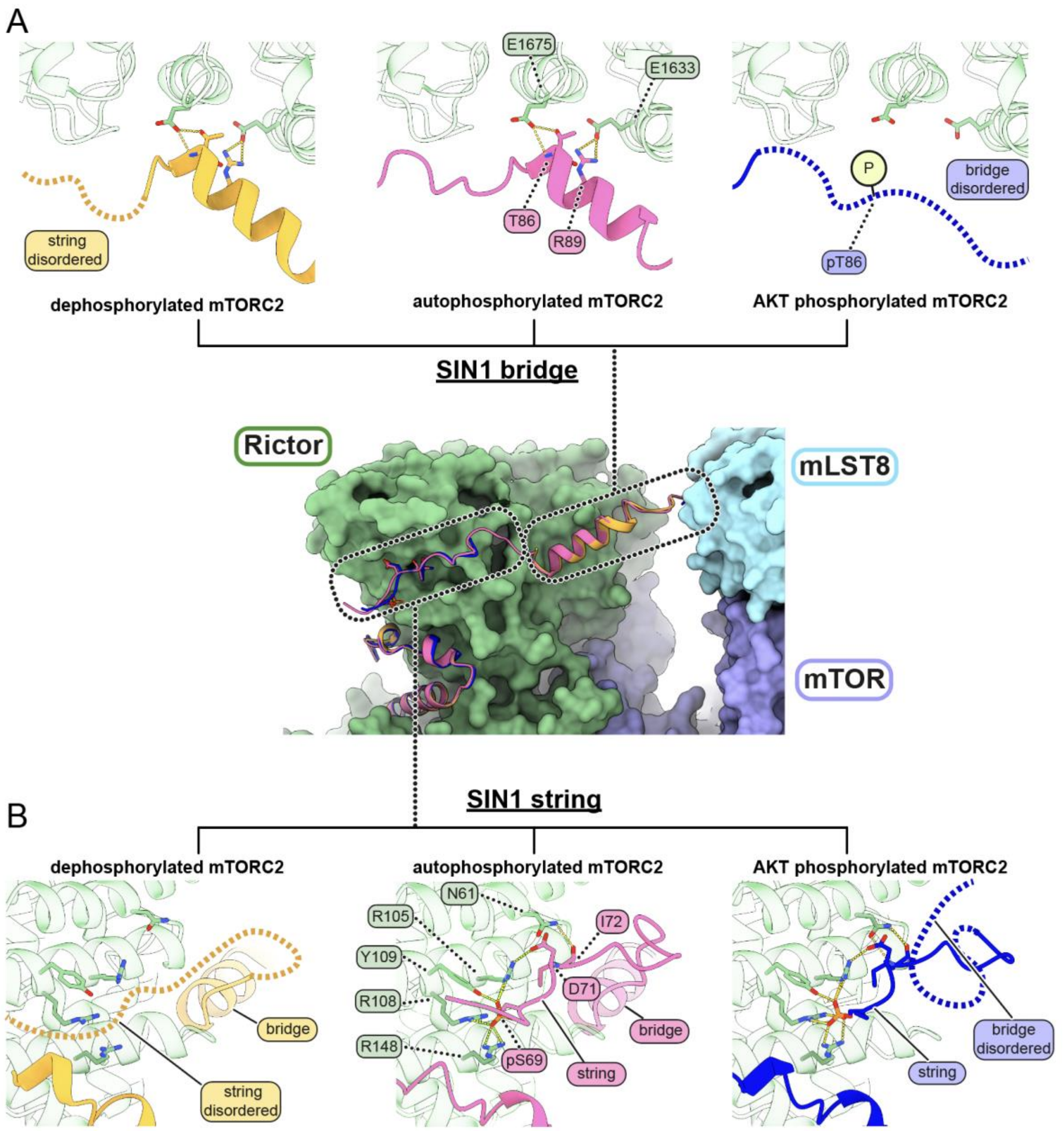

**Fig. S10. SIN1 bridge and string organization across all mTORC2 phosphorylation states.**

The middle panel shows an aligned overview of SIN1 subunits from dephospho- (orange), auto- (pink) and phospho- (blue) mTORC2 models. mTOR, mLST8 and Rictor are from the phospho-mTORC2 model and are shown as molecular surfaces. The top inset (A) shows the different states of the SIN1 bridge across mTORC2 phosphorylation. Both dephospho- and auto-mTORC2 have an ordered SIN1 bridge, stabilized by interaction of SIN1-T86 with Rictor-E1675 and SIN1-R89 with Rictor-E1633. In AKT-phosphorylated mTORC2 the bridge is disordered, driven by phosphorylation of SIN1-T86. The bottom inset (B) shows the different states of the SIN1 string across mTORC2 phosphorylation. Both auto- and AKT-phospho-mTORC2 have an ordered SIN1-string, stabilized by interaction of the autophosphorylated SIN1-S69 with Rictor-R105, R108, Y109 and R148. Dephospho-mTORC2 lacks an ordered string because of the absence of SIN1-S69 autophosphorylation. In all panels, key residues are shown as sticks, these common residues are labelled for auto-mTORC2 in the middle panel.

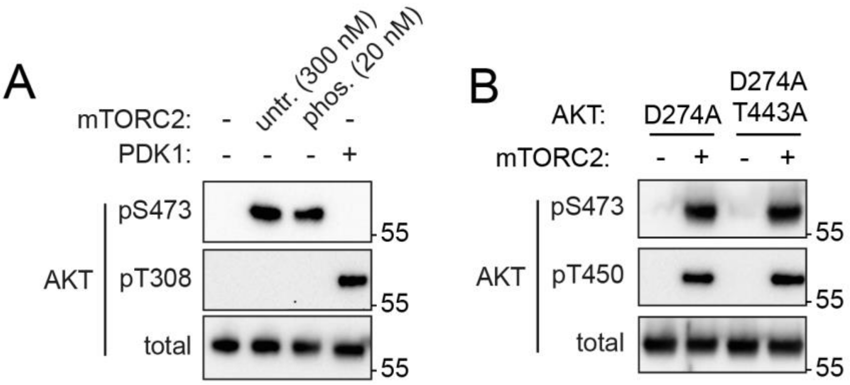

**Fig. S11. AKT activity is not required for S473 phosphorylation.**

(A) mTORC2 can phosphorylate pS473 within kinase dead AKT. Western blotting of pT308 and pS473 phosphorylation of 25 µM $AKT^{D274A}$ after treatment with untreated mTORC2 (untr., 300 nM), AKT-phospho-mTORC2 (phos., 20 nM) or PDK1 (300 nM) for 10 min at 30°C.

(B) Western blotting of T450 and S473 phosphorylation of either 25 µM $AKT^{D274A}$ or $AKT^{D247A/T443A}$ after treatment with 0.3 µM Untr-mTORC2 for 20 min at 30°C.

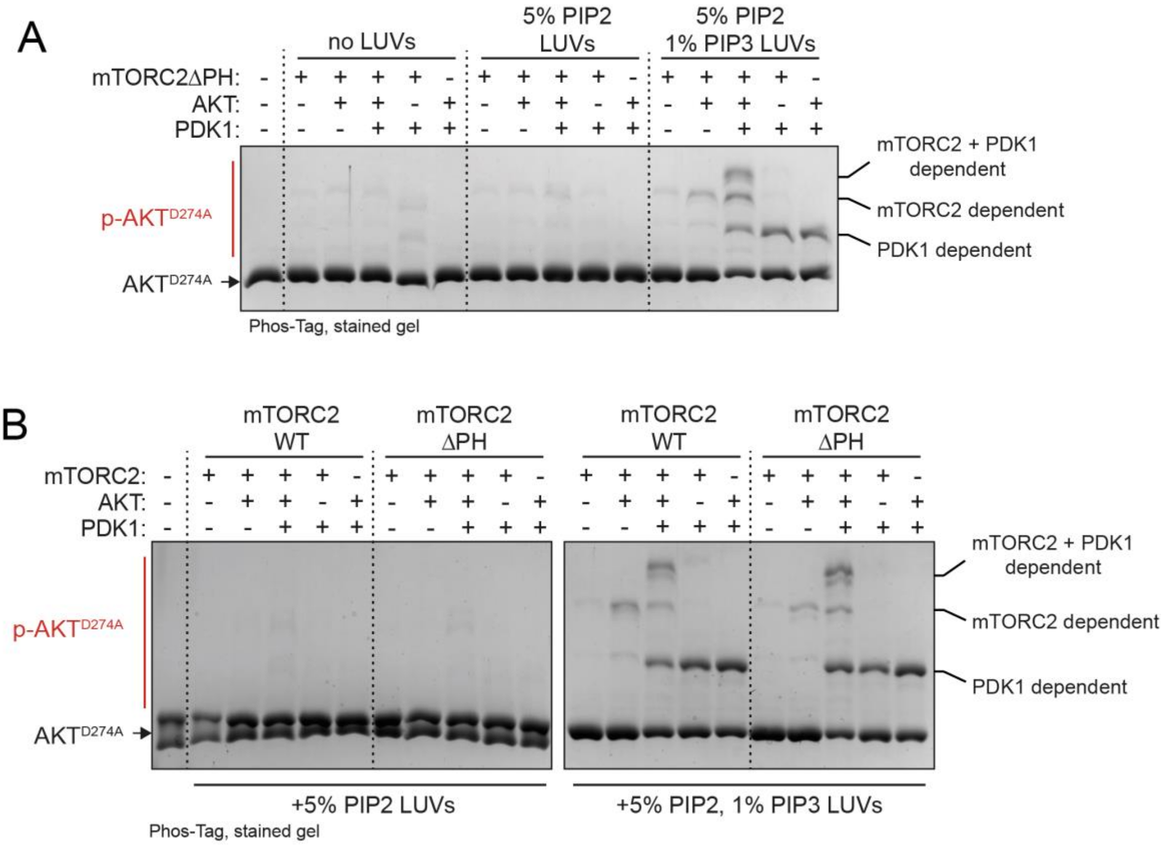

**Fig. S12. Analysis of mTORC2/AKT positive feedback with SIN1ΔPH.**

(A) mTORC2 containing SIN1 PH domain deletion (mTORC2ΔPH) is activated on membranes by AKT/PDK1. The indicated combinations of dephosphorylated enzymes (all 30 nM) and liposomes (0.4 mg/mL) were incubated in mTORC2 activity assays with 25 μM AKT$^{D274A}$ substrate for 10 min at 30°C.

(B) Activation of mTORC2 is unaffected by SIN1-PH deletion. The indicated combinations of dephosphorylated enzymes (all 30 nM) and liposomes (0.4 mg/mL) were incubated in mTORC2 activity assays with 25 μM AKT$^{D274A}$ substrate for 10 min at 30°C.

In both gels, the lowest phospho-AKT band (marked "PDK1-dependent") is PDK1 driven T308 phosphorylation, as this band is only observed in the presence of PDK1 and is stimulated by PIP3. The middle band (mTORC2-dependent) is mTORC2 phosphorylation of AKT as observed in other assays in this work and which is only generated in the presence of mTORC2. The top band (mTORC2+PDK1-dependent) is likely mTORC2-phosphorylated AKT with further T308 phosphorylation by PDK1 and is only generated in the presence of both mTORC2 and PDK1. Therefore, the total mTORC2 activity is represented by the sum of the top 2 bands.

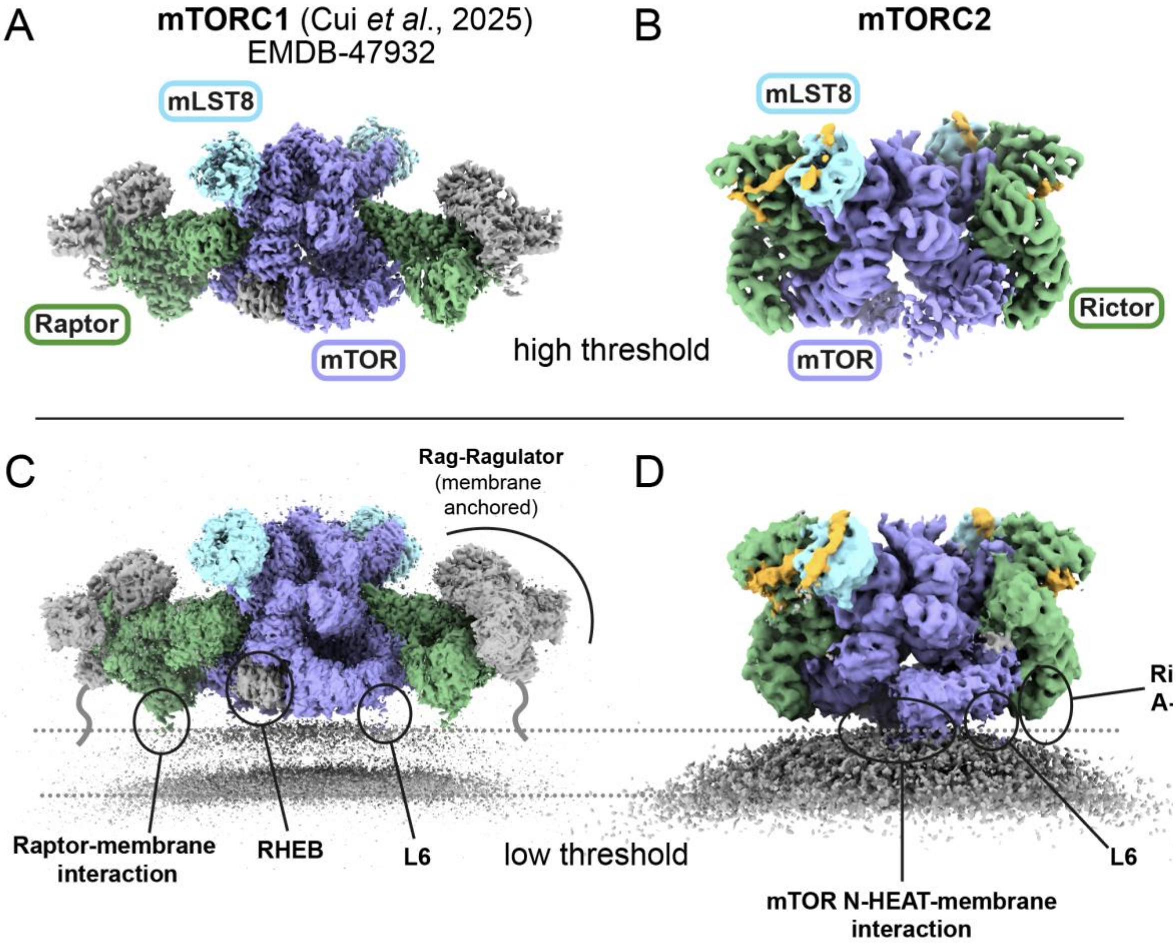

**Fig. S13. Comparison of mTORC1 and mTORC2 membrane binding.** Cryo-EM density maps are shown for membrane-bound mTORC1-Rag-Ragulator-RHEB (EMD-47932) (*38*) and mTORC2 (this study). Maps are shown at both high (A and B) and low (C and D) threshold levels to aid in visualization of the membrane and low-resolution features. The mTORC2 N-HEAT interacts with the membrane (highlighted in D), while mTORC1 appears to float above the membrane. This could be due to Raptor membrane interaction acting as a spacer, as well as binding of the mTORC1 N-HEAT binding to RHEB (both highlighted in C). The Raptor interaction area is positioned similarly to the A-site in Rictor (highlighted in D). Loop L6, which Cui *et al*. showed to be critical for membrane activation of mTORC1 and we also observe to be membrane proximal in mTORC2, is highlighted in C and D.

**Table S1. LUV lipid mixtures used in this study.**

| Composition | Name in Figures |
|---|---|
| 35% DOPE, 30% DOPC, 15% DOPS, 20% cholesterol | No PIP |
| 35% DOPE, **25% DOPC**, 15% DOPS, 20% cholesterol, **5% DOPIP2** | 5% PIP2 |
| 35% DOPE, **25% DOPC**, 15% DOPS, 20% cholesterol, **5% DOPIP3** | 5% PIP3 |
| 35% DOPE, **24% DOPC**, 15% DOPS, 20% cholesterol, **5% DOPIP2**, **1% DOPIP3** | 5% PIP2 1% PIP3 |

**Table S2. Cryo-ET data collection, refinement and validation statistics for membrane-bound mTORC2 subtomogram averaging**

| | |
|---|---|
| **Data collection and processing** | PDB 9TPW<br>EMD-56117 |
| Magnification | 81,000 |
| Voltage (kV) | 300 |
| Electron exposure (e–/Å$^2$) | 3.4 (per tilt image)<br>136 (per tilt-series) |
| Defocus range (μm) | -2.8 to -3.8 |
| Pixel size (Å) | 1.514 |
| Symmetry imposed | C1 |
| Initial particle images (no.) | 160,332 |
| Final particle images (no.) | 38,469 |
| Map resolution (Å) | 6.42 |
| FSC threshold | (0.143) |
| Map resolution range (Å) | 5.67 – 11.45 |
| **Refinement** | |
| Initial model used (PDB code) | 9T92 (this study), AlphaFold |
| Model resolution (Å) | 6.74 |
| FSC threshold | (0.5) |
| Model composition | |
| Non-hydrogen atoms | 61,294 |
| Protein residues | 7,634 |
| Ligands | 4 |
| *B* factors (Å$^2$) | |
| Protein | 181.43 |
| Ligand | 145.94 |
| R.m.s. deviations | |
| Bond lengths (Å) | 0.0076 |
| Bond angles (°) | 1.37 |
| Validation | |
| MolProbity score | 1.14 |
| Clashscore | 2.27 |
| Poor rotamers (%) | 0.00 |
| Ramachandran plot | |
| Favored (%) | 97.28 |
| Allowed (%) | 2.69 |
| Disallowed (%) | 0.03 |

**Table S3. Cryo-EM data collection, refinement and validation statistics for mTORC2 phosphorylation states**

| | Dephosphorylated mTORC2 | | Autophosphorylated mTORC2 | | AKT-phosphorylated mTORC2 | |
|---|---|---|---|---|---|---|
| **Data collection and processing** | **Dimer** | **Focused protomer** | **Dimer** | **Focused protomer** | **Dimer** | **Focused protomer** |
| | PDB 9TDS EMD-55803 | PDB 9TDT EMD-55804 | PDB 9T92 EMD-55714 | PDB 9T93 EMD-55715 | PDB 9T7J EMD-55637 | PDB 9T94 EMD-55716 |
| Magnification | 130,000 | | 130,000 | | 130,000 | |
| Voltage (kV) | 300 | | 300 | | 300 | |
| Electron exposure (e–/Å$^2$) | 45 | | 50 | | 50 | |
| Defocus range (μm) | -0.8 to -2.8 | | -0.8 to -2.8 | | -0.8 to -2.8 | |
| Pixel size (Å) | 0.955 | | 0.955 | | 0.955 | |
| Symmetry imposed | C1 | | C1 | | C1 | |
| Initial particle images (no.) | 1,543,822 | | 1,861,378 | | 1,126,327 | |
| Final particle images (no.) | 191,304 | | 143,361 | | 210,421 | |
| Map resolution (Å) | 3.30 | 2.98 | 3.13 | 2.84 | 3.00 | 2.63 |
| FSC threshold | (0.143) | (0.143) | (0.143) | (0.143) | (0.143) | (0.143) |
| Map resolution range (Å) | 2.94 – 11.20 | 2.80 – 6.13 | 2.75 – 11.40 | 2.68 – 5.58 | 2.60 – 10.05 | 2.41 – 5.75 |
| **Refinement** | **Dimer** | **Focused protomer** | **Dimer** | **Focused protomer** | **Dimer** | **Focused protomer** |
| Initial model used | 9T92 (this study) | 9T93 (this study) | 9T92 (this study) | ModelAngelo, AlphaFold | 9T92 (this study) | 9T93 (this study) |
| Model resolution (Å) | 3.38 | 3.08 | 3.26 | 2.97 | 3.05 | 2.70 |
| FSC threshold | (0.5) | (0.5) | (0.5) | (0.5) | (0.5) | (0.5) |
| Model composition | | | | | | |
| Non-hydrogen | 52,832 | 25,046 | 53,168 | 25205 | 52,802 | 24,959 |
| atoms | 6,562 | 3,107 | 6,601 | 3126 | 6,548 | 3,090 |
| Protein residues | 6 | 3 | 6 | 6 | 10 | 5 |
| Ligands | | | | | | |
| *B* factors (Å$^2$) | | | | | | |
| Protein | 102.75 | 64.89 | 130.16 | 75.52 | 87.24 | 67.10 |
| Ligand | 138.37 | 83.56 | 143.95 | 108.7 | 109.19 | 84.06 |
| R.m.s. deviations | | | | | | |
| Bond lengths (Å) | 0.0079 | 0.0036 | 0.0101 | 0.0038 | 0.0043 | 0.0039 |
| Bond angles (°) | 1.16 | 0.99 | 1.23 | 1.06 | 1.04 | 1.02 |
| Validation | | | | | | |
| MolProbity score | 0.93 | 0.77 | 1.01 | 0.72 | 0.84 | 0.88 |
| Clashscore | 1.26 | 0.54 | 1.81 | 0.59 | 0.81 | 1.00 |
| Poor rotamers (%) | 0.00 | 0.00 | 0.00 | 0.00 | 0.00 | 0.0 |
| Ramachandran plot | | | | | | |
| Favored (%) | 97.59 | 97.59 | 97.56 | 97.93 | 97.57 | 97.57 |
| Allowed (%) | 2.41 | 2.41 | 2.44 | 2.07 | 2.43 | 2.40 |
| Disallowed (%) | 0.00 | 0.00 | 0.00 | 0 | 0.00 | 0.03 |

**Table S4. Estimation of physiological enzyme concentrations.** Cellular concentrations for mTORC2 components, AKT and PDK1, taken from the HeLa Spatial Proteome (HSP; http://mapofthecell.biochem.mpg.de/, (*70*)), PaxDB (https://pax-db.org/,(*71*)) and OpenCell (https://opencell.sf.czbiohub.org/, (*72*)).
SIN1 was used for consideration of mTORC2 concentration, as it appears to be the limiting subunit.
For PaxDB estimates, reported parts per million values (ppm) were converted to nM using $C = (k \times A)/N_A$, where $k$ =~3 x $10^6$ proteins/fL, A = abundance in ppm and $N_A$ is Avogadro's number (*73*).

| | **Reported cellular concentration (nM):** | | |
|---|---|---|---|
| **Database:** | **HSP** | **PaxDB** | **OpenCell** |
| mTOR | 50 | 50 | 110 |
| mLST8 | 80 | 70 | 430 |
| Rictor | 38 | 25 | 27 |
| **SIN1** | **10** | **25** | **28** |
| **AKT1** | **37** | **125** | **37** |
| **PDK1** | **13** | **20** | **2** |
